# Feature-space selection with banded ridge regression

**DOI:** 10.1101/2022.05.05.490831

**Authors:** Tom Dupré la Tour, Michael Eickenberg, Anwar O. Nunez-Elizalde, Jack L. Gallant

## Abstract

Encoding models provide a powerful framework to identify the information represented in brain recordings. In this framework, a stimulus representation is expressed within a feature space and is used in a regularized linear regression to predict brain activity. To account for a potential complementarity of different feature spaces, a joint model is fit on multiple feature spaces simultaneously. To adapt regularization strength to each feature space, ridge regression is extended to banded ridge regression, which optimizes a different regularization hyperparameter per feature space. The present paper proposes a method to decompose over feature spaces the variance explained by a banded ridge regression model. It also describes how banded ridge regression performs a feature-space selection, effectively ignoring non-predictive and redundant feature spaces. This feature-space selection leads to better prediction accuracy and to better interpretability. Banded ridge regression is then mathematically linked to a number of other regression methods with similar feature-space selection mechanisms. Finally, several methods are proposed to address the computational challenge of fitting banded ridge regressions on large numbers of voxels and feature spaces. All implementations are released in an open-source Python package called Himalaya.

## Introduction

A fundamental problem in neuroscience is to identify the information represented in different brain areas. In the encoding model framework, this identification amounts to finding the features of the stimulus (or task) that predict the activity in each brain area [Wu et al., 2006; Naselaris et al., 2011]. Encoding models have been used extensively to predict blood oxygen level dependent (BOLD) signals in functional magnetic resonance imaging (fMRI). When encoding models are used independently to model each spatial sample in fMRI recordings (*i*.*e*. on each *voxel*), they are called *voxelwise* encoding models. Voxelwise encoding models have been used to predict BOLD activity generated by visual images and videos [Hansen et al., 2004; Kay et al., 2008; Nishimoto et al., 2011; Huth et al., 2012; Eickenberg et al., 2016; Wen et al., 2018; Lescroart and Gallant, 2019], music [Kell et al., 2018], semantic concepts [Mitchell et al., 2008; Wehbe et al., 2014; Huth et al., 2016] and narrative language [de Heer et al., 2017; Huth et al., 2016; Jain and Huth, 2018; Deniz et al., 2019]. Encoding models have also been applied to data acquired using other neuroimaging techniques, such as neurophysiological recordings (see [Wu et al., 2006] and references therein), calcium imaging [Miri et al., 2011; Pinto and Dan, 2015; Oldfield et al., 2020], magneto-encephalography [Schwartz et al., 2019; Yang et al., 2019], and electro-corticography [Holdgraf et al., 2016, 2017].

In the encoding model framework, brain activity is recorded while subjects perceive a stimulus or perform a task. Then, a set of features (also known as a *feature space*) is extracted from the stimulus or task at each point in time. For example, a video might be represented in terms of amount of motion in each part of the screen [Nishimoto et al., 2011], or in terms of semantic categories of the objects and actions present in the scene [Huth et al., 2012]. Each feature space corresponds to a different representation of the stimulus- or task-related information. The encoding model framework aims to identify if each representation is similarly encoded in brain activity. Each feature space thus corresponds to a hypothesis about the stimulus- or task-related information encoded in brain activity. To test the hypothesis associated with a feature space, a regression model is trained to predict brain activity from the feature space. If the regression model predicts brain activity significantly in a voxel, then one may conclude that the information represented in the feature space is also represented in brain activity [Wu et al., 2006; Naselaris et al., 2011] (see more details about encoding models in Section 1).

Comparing the prediction accuracy of different feature spaces amounts to comparing competing hypotheses. In each brain voxel, the best-predicting feature space corresponds to the best hypothesis about the information encoded in this voxel. However, it is important not to view feature-space comparison as a winner-take-all situation. It is possible (and in fact common) to find that a brain area represents multiple feature spaces simultaneously. To take this possibility into account, a joint regression is fit on multiple feature spaces.

The joint regression automatically combines the information from all feature spaces to maximize the joint prediction accuracy. Then, a variance decomposition method (*e*.*g*. variance partitioning [Lescroart et al., 2015; Çukur et al., 2016; de Heer et al., 2017; Lescroart and Gallant, 2019; LeBel et al., 2021]) is used to decompose the variance explained by the joint model into separate contributions from each feature space. However, variance partitioning is limited in practice to small numbers of feature spaces *m* (*e*.*g*. 2 or 3), because it decomposes the variance into 2^*m*^ − 1 values that are hard to interpret. There is a need for a method to decompose the variance over larger numbers of feature spaces.

The regression model trained to predict brain activity from a feature space is usually regularized in order to improve generalization. The most common regularized regression model is ridge regression (see more details in Section 1.2). However, ridge regression is not suited to fit joint models on multiple feature spaces, because it uses the same regularization hyperparameter for all feature spaces. In practice, it is rare for all feature spaces to require the same level of regularization, because the optimal level of regularization depends on factors such as the number of features, the feature covariances, and the predictive power of the feature space. To address this challenge, ridge regression is naturally extended to use a separate regularization level per feature space. This extension is called *banded ridge regression* [Nunez-Elizalde et al., 2019]. By learning a separate regularization hyperparameter for every feature space, banded ridge regression can optimize regularization strength separately for each feature space. As a side effect, learning separate hyperparameters leads to a feature-space selection mechanism that was not described in the original paper [Nunez-Elizalde et al., 2019]. There is a need for a proper description of this mechanism.

Another challenge with banded ridge regression is its computational cost. Because this method requires the optimization of a separate hyperparameter for every feature space, the optimization algorithm is necessarily more complicated than that used for simple ridge regression. For example, using a grid search (as is common in ridge regression) is impractical in banded ridge regression with more than three or four feature spaces because its computational cost scales exponentially with the number of feature spaces. This computational challenge is particularly difficult when modeling hundreds of thousands of voxels independently. There is a need for an algorithm to efficiently fit models with banded ridge regression.

### Overview of the paper

This work expands on earlier banded ridge regression work from our laboratory [Nunez-Elizalde et al., 2019] in three ways, described respectively in the three sections.

Section 1 first provides a brief overview of the encoding model framework. Then, it proposes an alternative method to variance partitioning, to decompose variance over large numbers of feature spaces. The proposed method is based on the product measure [Hoffman, 1960; Pratt, 1987], whose computational cost scales linearly with the number of feature spaces. This variance decomposition can be applied to any linear regression fitting method, including ridge regression and banded ridge regression.

Section 2 reviews banded ridge regression, and describes how it performs a feature-space selection by discarding non-predictive or redundant feature spaces. This feature-space selection mechanism explains how banded ridge regression is able to effectively disentangle correlated feature spaces [Nunez-Elizalde et al., 2019]. Then, it discusses the relationship between banded ridge regression and other regression methods with a similar feature-space selection mechanism, such as automatic relevance determination, multiple-kernel learning, and the group lasso.

Section 3 addresses the practical challenge of solving banded ridge regressions with many feature spaces on large numbers of voxels. Two methods are proposed to efficiently solve banded ridge regression. The first uses hyperparameter random search over a Dirichlet distribution, while the second uses hyperparameter gradient descent with implicit differentiation. Both methods provide speed improvements of several orders of magnitude compared to the Bayesian search proposed earlier [Nunez-Elizalde et al., 2019]. Finally, all implementations are made available in an open-source Python package called *Himalaya*^1^, with both CPU and GPU support, rigorous unit-testing, and extensive documentation.

This paper focuses on fMRI datasets and the associated terminology. However, the methods presented here can also be applied to data acquired using other neuroimaging modalities, and more generally to any similarly structured regression setting. The methods could even be used to integrate across multiple neuroimaging modalities Rasero et al. [2021].

## 1 Variance decomposition over feature spaces

This section starts with a brief overview of the encoding model framework, introducing key concepts such as linear regression, ridge regression, explained variance, and feature-space comparison. Then, it explains the benefit of fitting joint models over multiple feature spaces, and discusses how to decompose the explained variance over feature spaces by adapting the product measure [Hoffman, 1960; Pratt, 1987].

### 1.1 Linear regression

The regression method used in encoding models is often linear in the parameters. Using linear regression has several benefits compared to non-linear regression. The main benefit is to only focus on *explicit* representations [Kamitani and Tong, 2005; Wu et al., 2006; Naselaris et al., 2011; Kriegeskorte and Kievit, 2013; King et al., 2018; Ivanova et al., 2021]. With non-linear regression, some features might be able to predict brain activity even though they do not explicitly encode the same information as the brain region (*e*.*g*. pixel features do not explicitly encode the semantic content of an image, even though semantic content can be extracted from the pixels). Restricting the regression method to be linear is thus a way to only consider features that explicitly represent brain activity. Additionally, linearity enables the use of various methods to interpret the regression model, such as variance partitioning or feature importance methods (see Section 1.5). Finally, linear regression methods are computationally efficient and robust to small sample sizes that are common in neuroimaging studies. Note that encoding models that use linear regression are sometimes called *linearized* encoding models [Wu et al., 2006; Naselaris et al., 2011], to emphasize that feature extraction can be non-linear.

Mathematically, let us note *X* ∈ ℝ^*n*×*p*^ the matrix of features, with *n* samples and *p* features, and *y* ∈ R^*n*^ the brain activity vector (or target vector). Linear regression seeks a weight vector *b* ∈ ℝ^*p*^ such that the target vector is approximated as a linear combination of the features *y* ≈ *Xb*. It is standard to model the error as additive Gaussian white noise, so that *y* = *Xb* + *ε*, with a noise vector *ε* ∈ R^*n*^ drawn from a centered Gaussian distribution. (The Gaussian assumption is reasonable for de-trended fMRI recordings, but might not be valid for single-trial neurophysiology). This regression method is called *ordinary least squares*, and can be written as an optimization problem

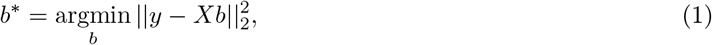

where 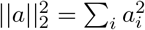 is the squared *L*_2_ norm.

The main limitation of ordinary least squares is that it is unstable in some situations. To understand this issue, note that solving ordinary least squares amounts to inverting the eigenvalues *σ*_*i*_ of the matrix *X*^⊤^*X*. When features are correlated, some eigenvalues can be small, which reduces the stability of the inversion (a small change in the features can have a large effect on the result). In some cases, some eigenvalues can even be equal to zero, which makes the inversion ill-defined. Eigenvalues equal to zero can arise when some features are fully redundant (*e*.*g*. one feature being the sum of two other features), or when the number of independent features is greater than the number of samples (p > n). The system is then called underdetermined, and the optimization problem has infinitely many solutions, of which most do not generalize well to new data. To solve these issues, ordinary least squares can be extended into ridge regression [Hoerl and Kennard, 1970].

### 1.2 Ridge regression

Ridge regression [Hoerl and Kennard, 1970] is a generalization of ordinary least squares which adds a regularization term to the optimization problem. The optimization problem becomes

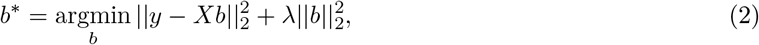

where *λ* > 0 is a hyperparameter controlling the regularization strength. This regularization improves the robustness of the model by adding a positive value *λ* to all eigenvalues *σ*_*i*_ ≥ 0 of *X*^⊤^*X* before the matrix inversion. Inverting (*σ*_*i*_ + *λ*) instead of *σ*_*i*_ reduces the instability caused by small eigenvalues, and allows inverting underdetermined systems (where *σ*_*i*_ = 0). Small eigenvalues and underdetermined systems are common when using large numbers of features. Because naturalistic stimuli often involve large numbers of highly correlated features, this regularization scheme facilitates the analysis of naturalistic experiments and so increases ecological validity [Wu et al., 2006; Huth et al., 2016; Sonkusare et al., 2019].

In ridge regression, the regularization hyperparameter *λ* must be estimated from the data. If the estimated value is too low then the model will tend to overfit to noise in the data, and if it is too high then the model will not predict as well as it might. The regularization parameter is typically selected through a grid search with cross-validation [Hastie et al., 2009, Ch. 7]. First, a set of hyperparameter candidates *λ* is defined, typically over a grid of logarithmically spaced values to span multiple orders of magnitude. Then, the data set is split into a training set (*X*_train_, *y*_train_) and a validation set (*X*_val_, *y*_val_). The regression model is fit on the training set, and its prediction accuracy is evaluated on the validation set. This process is performed for all hyperparameter candidates, and the candidate leading to the highest prediction accuracy is selected. To improve the robustness of the hyperparameter selection, the prediction accuracy is usually averaged over multiple splits of the data into a training and a validation set. Finally, a ridge regression is fit on the full data set using the selected hyperparameter.

Ridge regression is the most popular regression method in voxelwise encoding models. Other popular linear regression methods include the lasso [Tibshirani, 1996] or linear support-vector machines [Cortes and Vapnik, 1995]. However, both methods have a much larger computational cost than ridge regression. In particular, their computational cost increases linearly with the number of voxels. Because a typical fMRI dataset contains about 10^5^ voxels, fitting an independent regression model on each voxel is computationally challenging. In the case of ridge regression, the computational bottleneck (inverting the linear system) can be shared for all voxels (see Section 3.1 for more details). This is not the case in the lasso or in support-vector machines. For these reasons, solving ridge regression on large numbers of voxels is fast compared to solving the lasso or linear support-vector machines.

### 1.3 Explained variance

To estimate the prediction accuracy of a model, the model prediction is compared with the recorded brain response. However, encoding models that are relatively more complex are more likely to fit noise in the training data and so they may not generalize well to new data. To avoid artificially inflated performances due to overfitting, this comparison should be performed on a separate data set not used during model training nor during hyperparameter optimization. The ability to evaluate a model on a separate data set is critical to the encoding model framework. It provides a principled way to build complex models while controlling the amount of overfitting. This separate data set is either called the *test set* or the *validation set* depending on the community. This paper uses the term *test set*.

The two standard metrics to quantify prediction accuracy of linear regression models are the *R*^2^ score and the Pearson correlation *r*. This paper only uses the *R*^2^ score, due to the possibility to decompose the *R*^2^ score over feature spaces (see Section 1.5). The *R*^2^ score is generally defined as

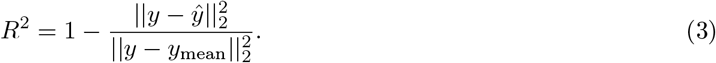

The *R*^2^ score is usually interpreted as the explained variance, *i*.*e*. the part of the variance that is explained by the model. This interpretation originates from ordinary least squares, where the prediction vector *ŷ* and the error vector *y* − *ŷ* are always orthogonal on the train set. Indeed, *ŷ* is the vector that minimizes the error 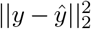 so it is the orthogonal projection of *y* over the subspace of all linear combinations of *X*. Because of this orthogonality, the variance of the brain activity vector *y* on the train set can be decomposed between the variance of the prediction vector and the variance of the error vector

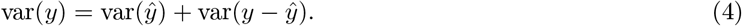

Because (*y* − *ŷ*) is zero-mean in ordinary least square on the train set, we get 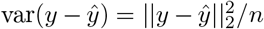 Then, dividing (4) by var(*y*) leads to

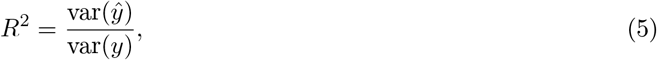

which is naturally interpreted as the explained variance ratio. However, in the test set and in the case of regularized regression methods such as ridge regression, the prediction vector *ŷ* and the error vector *y* − *ŷ* are not always orthogonal. Therefore, equations (3) and (5) are not equivalent, and only (5) corresponds to the explained variance ratio. The definition in (3) is however preferred because it is more general [Kvålseth, 1985], and it is still (improperly) interpreted as the explained variance.

The *R*^2^ score takes values in (−∞, 1]. Larger values are better, and a value of 1 corresponds to a perfect prediction. A value of 0 corresponds to the baseline prediction of a constant signal equal to the mean value of *y*. Negative values can be obtained when the variance of the error 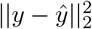 is larger than the variance of the signal 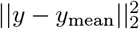 This is often (though not always) a sign of overfitting.

#### Noise floor

Accurate interpretation of the *R*^2^ score requires comparison to a lower bound given by the statistical significance threshold, and an upper bound given by the noise ceiling. The statistical significance threshold is often estimated by first randomly permuting the prediction and then computing the *R*^2^ score for each permuted prediction. This produces a distribution of *R*^2^ scores under the null hypothesis that the predictions are independent of the recorded brain activity. The statistical significance threshold can be identified from this distribution for any critical p-value that the experimenter might choose. Because this threshold reflects what would be expected to be obtained by chance alone, it is also called the *noise floor*.

Importantly, a statistically significant value does not necessarily correspond to a strong effect size. Any small effect can be significant if the dataset is large enough. Therefore, the effect size should always be reported along with the significance threshold. The *R*^2^ score gives an intuitive quantification of the effect size as the ratio of variance explained by the model.

#### Noise ceiling

Second, it is necessary to estimate an upper bound for the *R*^2^ score. In theory the *R*^2^ score can reach a value as high as 1, but this is never achieved in practice when there is noise in the data. Therefore, interpretation of the *R*^2^ score also requires estimating the upper bound on possible values of *R*^2^, the *noise ceiling*. If brain responses are stationary with respect to repetitions of the experiment then this upper bound can be estimated by repeating the same experiment several times. Given stationarity, some fraction of the brain response will be identically reproduced over repetitions (the *reproducible signal*), while the rest will vary across repetitions (the *noise*). Because an encoding model can only explain reproducible signals and not noise, the variance of the reproducible signal can be used to estimate the variance that could be potentially explained by an ideal encoding model. The explainable variance can then be used to compute the maximum *R*^2^ score that can be obtained with an encoding model [Sahani and Linden, 2003; Hsu et al., 2004; Schoppe et al., 2016].

### 1.4 Feature-space comparison

A key element of the encoding model framework is the ability to compare different feature spaces based on their prediction accuracies. A feature space is a set of features that reflects information contained in the experimental stimulus or task. For example, stimulus-related features might be obtained by computing the spatio-temporal wavelet transform of images and videos [Kay et al., 2008; Nishimoto et al., 2011], by labeling the actions and objects present in visual scenes [Huth et al., 2012], or by computing word embeddings that capture the semantic properties of speech [Wehbe et al., 2014; Huth et al., 2016; Deniz et al., 2019]. Task-related features might be defined as indicator variables of attention tasks [Çukur et al., 2013], behavioral states of a video game task [Zhang et al., 2021], or progress during a navigation task [Zhang and Gallant, 2020].

In principle a feature space could reflect any type of stimulus- or task-related information that might potentially be represented in the brain. Thus, a feature space can be viewed as a specific scientific hypothesis about functional brain representations. To test the hypothesis associated with some specific feature space, a regression model is trained to predict brain activity from that feature space. When the regression model has significant prediction accuracy in a particular brain voxel, it suggests that the information represented in the features correlates with the functional activity measured in that brain voxel [Wu et al., 2006; Naselaris et al., 2011].

Multiple hypotheses can be tested on the same dataset by instantiating multiple feature spaces and then comparing their prediction accuracies. If the prediction accuracy of one feature space is significantly higher than the performance of other feature spaces, then it is common to conclude that the corresponding hypothesis provides the best description about how information is represented in the voxel [Nishimoto et al., 2011; Çukur et al., 2016; Lescroart et al., 2015; de Heer et al., 2017]. However, this approach ignores the possibility that a feature space with lower performance might reflect some information that is represented in the brain, but which is not captured by the best predicting feature space. In this case, both feature spaces would be complementary, in the sense that using both of them jointly would provide a better description of brain function than would be obtained by using either one individually.

By fitting a joint model using multiple feature spaces simultaneously, it becomes possible to optimize the use of each feature space within the model. A joint model maximizes the model prediction accuracy, taking into account potential complementarity of feature spaces. One can then make inferences about the information encoded in a given voxel by decomposing the prediction accuracy of the joint model over the feature spaces.

### 1.5 Variance decomposition

In a joint model, it is likely that all feature spaces will not contribute equally to the prediction. Some feature spaces might predict a large fraction of brain activity while other feature spaces might have almost no effect. To interpret a joint model, it is thus essential to quantify the relative importance of each feature space in the model. However, most work on this issue has focused on the importance of individual features, not feature spaces. Many *feature importance* measures have been proposed and discussed in the linear regression literature (see reviews in [Nathans et al., 2012; Grömping, 2015]). The most common approach is to standardize the features to zero mean and unit variance, and to then use the magnitude of the regression weights 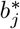 as a feature importance measure. Another commonly used feature importance measure is the correlation coefficient *r*(*X*_*j*_, *y*) between feature *X*_*j*_ and the target vector *y* (*e*.*g*. the activity of one voxel). Both measures quantify different aspects of feature importance, but they also have limitations (see [Grömping, 2015] for further discussion).

One limitation of both feature importance measures described above is that they are not variance decompositions. To be a variance decomposition, a measure must have two properties. First, the sum over the feature importance measure for all features must be equal to the *R*^2^ score of the model. Second, when all features are orthogonal then the importance of each feature must be equal to the *R*^2^ score obtained using each feature alone. With these two properties, a feature importance measure can be naturally summed over multiple features, to get the relative importance of entire feature spaces. Multiple feature importance measures have these two properties, and each has different limitations (see a list of properties and measures in [Grömping, 2015]). One variance decomposition measure is called variance partitioning, and has been used in the past in voxelwise encoding models. However, variance partitioning is impractical for more than 2 or 3 feature spaces (see Section 1.7). Therefore, this paper focuses on the product measure (described in the next section), because its computational cost scales linearly with the number of feature spaces.

### 1.6 The product measure

The product measure [Hoffman, 1960; Pratt, 1987] is a feature importance measure. It is defined as the product 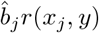 where 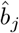 is the regression weight associated with feature *x*_*j*_, and *r*(*x*_*j*_, *y*) is the correlation between feature *x*_*j*_ and the target vector *y*. (This definition assumes that *y* and all features *x*_*j*_ are standardized to have zero mean and unit variance.) Importantly, when the linear regression is fit with ordinary least squares, the product measure is a variance decomposition on the training set. Indeed, when summing over features *j*, we have

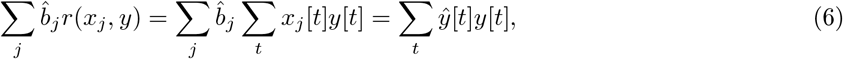

where *y*[*t*] is the value of *y* at time *t*. On the other hand, the *R*^2^ score can be written

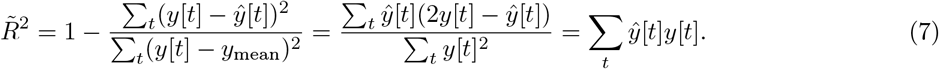

Here, the first equality uses the fact that *y* is zero-mean. The second equality uses the fact that with ordinary least squares and on the train set, the error vector *ŷ* − *y* is orthogonal to the target vector *y*, that is Σ_*t*_(*ŷ* [*t*] − *y*[*t*])*y*[*t*] = 0. Using both (6) and (7), the sum of the product measure over features is equal to the *R*^2^ score. Moreover, when all features *x*_*j*_ are orthogonal, the product measure decomposition is identical to using the *R*^2^ scores computed on each features alone. With these two properties the product measure is a variance decomposition, and it can be naturally summed over multiple features to obtain the relative importance of entire feature spaces.

Unfortunately, the variance decomposition property holds only for ordinary least squares and on the training set. Therefore, the product measure is not a variance decomposition when using a regularized regression method or when computing the *R*^2^ score on a held-out test set. To fix this issue, we propose here another definition of the product measure. If *y* is zero-mean, we can write

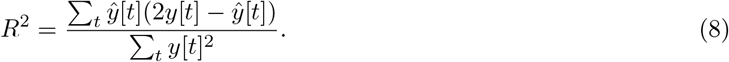

Then, we decompose the prediction *ŷ* into *ŷ* = Σ_*j*_ *ŷ*_*j*_, where *ŷ*_*j*_ = *X*_*j*_*b*_*j*_ is the sub-prediction computed on feature space *X*_*j*_ alone using the weights *b*_*j*_ of the joint model. From this decomposition, we define the variance explained by feature space *j* as

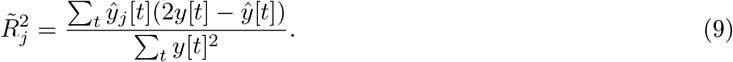

Using this new definition, the variance explained *R*^2^ by the joint model is the sum of the product measure over feature spaces: 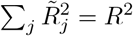 Moreover, when the sub-predictions *ŷ*_*j*_ are all orthogonal, the decomposition is identical to using the *R*^2^ scores computed on each sub-prediction 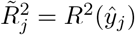 With these two properties the adapted product measure is a variance decomposition. Note that our new definition of the product measure is equal to the original definition in the case of ordinary least squares and on the training set. For this reason, this paper simply refers to it as the product measure.

One problem with the product measure is that it can yield negative values, even when the *R*^2^ score is positive [Thomas et al., 1998]. Negative values that are small relative to other values of the same decomposition might be ignored, but larger negative values are harder to interpret. Large negative values can occur when the regression model uses a positive weight on a feature negatively correlated with the target, or a negative weight on a feature positively correlated with the target. For example, given an orthonormal basis (*u, v*), two features *x*_1_ = *u* − *v* and *x*_2_ = −*u* + 2*v* can be used to predict *y* = *v* with a regression model *ŷ* = *x*_1_ + *x*_2_. The feature *x*_1_ is negatively correlated with the target, but it is used with a positive weight in the model. Thus, the sub-prediction *ŷ*_1_ = *x*_1_ is projected negatively on *ŷ*, which leads to a negative share 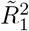 When considering the relative importance of entire feature spaces, this situation can sometimes be resolved by removing or merging feature spaces in the joint model. Note that this limitation of the product measure is also shared with many variance decomposition measures, including the variance partitioning measure described in the next subsection.

### 1.7 Other variance decomposition measures

Several other variance decomposition measures have been proposed in the linear regression literature, and in theory they could substitute for the product measure. One commonly used measure for variance decomposition of fMRI data is *variance partitioning* [Lescroart et al., 2015; Greene et al., 2016; Çukur et al., 2016; de Heer et al., 2017; Groen et al., 2018; Lescroart and Gallant, 2019; Snoek et al., 2019; LeBel et al., 2021]. This method is also known as commonality analysis in the statistics literature [Mood, 1969; Mayeske, 1969; Bring, 1995; Nathans et al., 2012]. Variance partitioning decomposes the explained variance into shared and unique components for each feature space, providing a more nuanced interpretation of the contribution of each feature space to predictions. However, variance partitioning decomposes the variance into 2^*m*^ − 1 values which are hard to interpret when the number of feature spaces *m* is large. To create these 2^*m*^ − 1 values, variance partitioning requires fitting a separate regression model on every subselection of feature spaces, which leads to 2^*m*^ − 1 regression models. The examples shown in this paper use up to 22 separate feature spaces. It would be impossible to use variance partitioning to understand the contributions of so many feature spaces. However, the product measure can still be used even in extreme cases such as this.

## 2 Banded ridge regression and feature-space selection

The previous section motivates the use of joint models fit over multiple feature spaces simultaneously, and proposes to use the product measure to decompose the variance explained by the model over feature spaces. This section tackles an additional challenge when fitting joint models, which is that different feature spaces might require different regularization strengths. This challenge can be solved using banded ridge regression [Nunez-Elizalde et al., 2019], which optimizes a separate regularization hyperparameter per feature space. Interestingly, this hyperparameter optimization leads to a feature-space selection mechanism. This section describes this feature-space selection mechanism, proposes a metric to quantify feature-space selection, and demonstrates its effect on two examples. Finally, a discussion presents the link between banded ridge regression and other regression methods with a similar feature-space selection mechanism.

### 2.1 A need for multiple regularization hyperparameters

When fitting a regularized regression such as ridge regression, the regularization strength has a large impact on the model prediction accuracy. When the regularization is too strong, then regression coefficients are too biased toward zero and so relevant features may be underused. On the other hand, when the regularization is too weak, then the model may overfit the training set and so generalization performance will be reduced. Therefore, the regularization strength must be optimized for the dataset under consideration. In particular, the optimal regularization strength depends on many characteristics of the feature space, such as its covariance, its number of features, or its predictive power. When fitting a joint model over multiple feature spaces simultaneously, ridge regression uses a single regularization hyperparameter and so it does not account for the different levels of regularization required for each feature space.

To solve this issue, ridge regression can be extended to *banded ridge regression* [Nunez-Elizalde et al., 2019], a special case of the more general Tikhonov regression [Tikhonov et al., 1977]. Banded ridge regression uses a separate regularization hyperparameter per feature space (Figure 2), and it learns these hyperparameters using cross-validation. Through these multiple hyperparameters, banded ridge regression is able to optimize the level of regularization for each feature space separately.

**Figure 1:**
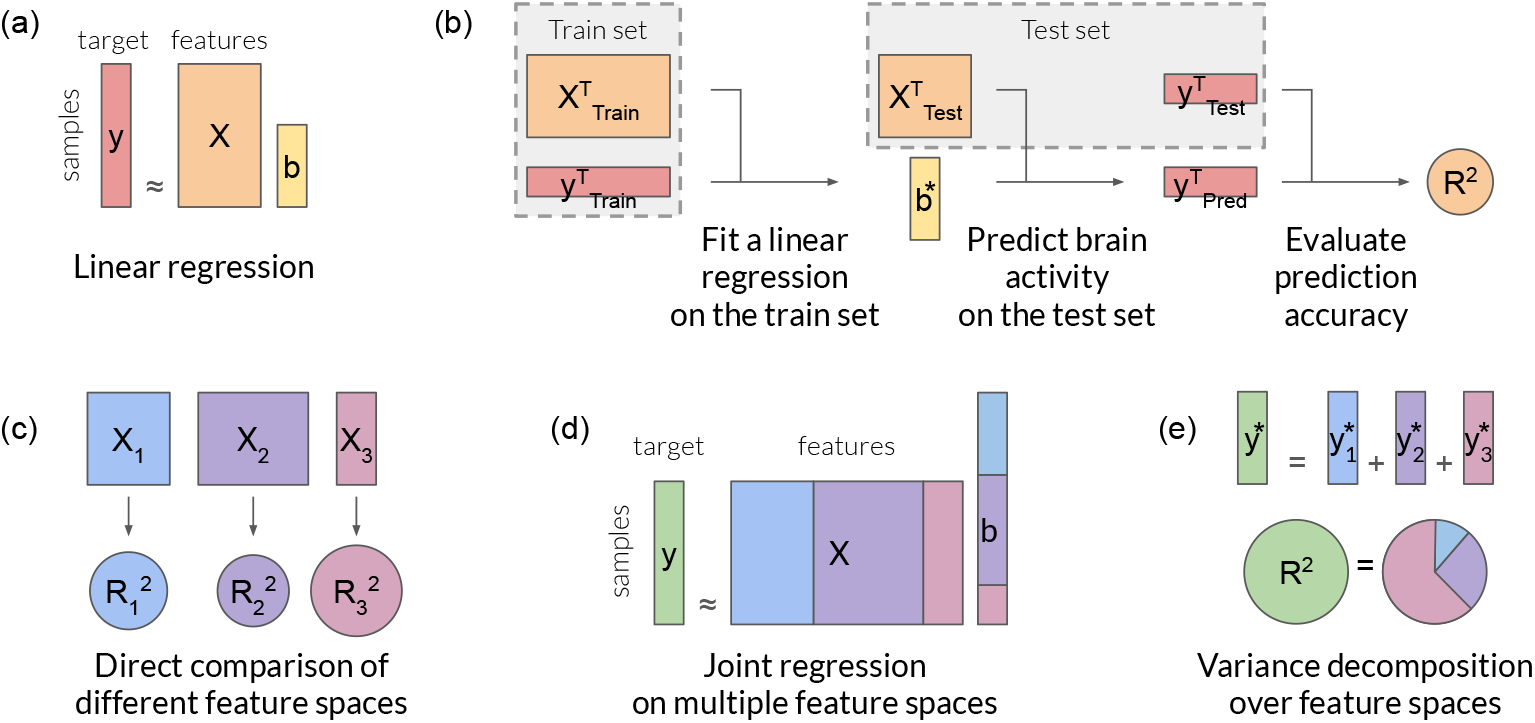
Overview of the encoding model framework. **(a)** The encoding model framework uses a regression model to predict a brain activity *y* ∈ ℝ^*n*^ with features *X* ∈ ℝ^*n×p*^ extracted from the stimulus (or task). The regression model is usually linear (see Section 1.1), *y* ≈ *Xb*, with a weight vector *b* ∈ ℝ^*p*^. Regularization can be also used to improve the regression model (see Section 1.2). **(b)** A cornerstone of the encoding model framework is the separation of the data into a train set and a test set. The regression model is fit on the train set only. Then, the fit model is used to predict brain activity on the test set. Finally, the prediction accuracy is evaluated on the test set by comparing the prediction vector with the recorded brain activity. The prediction accuracy is quantified for instance with the *R*^2^ score, which can be interpreted as the explained variance (see Section 1.3). **(c)** When multiple feature spaces are available, a separate model can be fit on each feature space, to compare the prediction accuracy of each model (see Section 1.4). However, considering only the best-predicting feature space ignores the possibility that feature spaces can be complementary. **(d)** To take into account complementarity between feature spaces, a joint regression can be fit on multiple feature spaces simultaneously. The feature spaces are concatenated into a large feature matrix *X*, and a single weight vector *b* is learned. **(e)** Then, the prediction vector *y* can be decomposed into a sum of partial predictions from each feature space *y*_*i*_ = *X*_*i*_*b*_*i*_. Finally, the explained variance *R*^2^ can be decomposed into the contribution of each feature space (see Section 1.5), for instance using the product measure (see Section 1.6) or variance partitioning (see Section 1.7).

**Figure 2:**
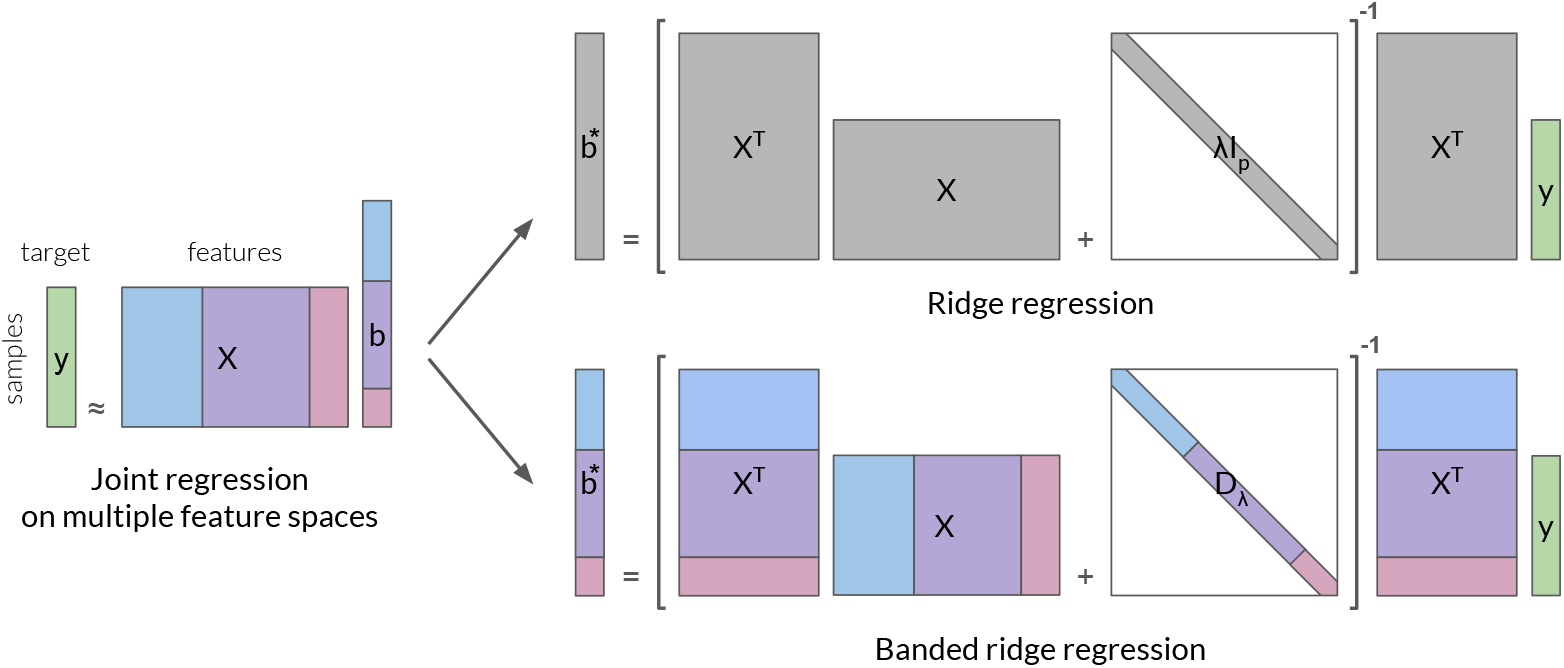
Ridge regression and banded ridge regression. In both regression methods, the brain activity vector *y* ∈ ℝ^*n*^ is approximated as a linear sum of the input features *X* ∈ ℝ^*n×p*^, using a weight vector *b* ∈ R^*p*^, specifically *y* ≈ *Xb*. In ridge regression, the weight vector is computed with *b*^*\**^ = (*X*^⊤^*X* + *λI*_*p*_)^−1^*X*^⊤^*y*, where *λ* > 0 is a regularization hyperparameter common to all features, and *I*_*p*_ ∈ ℝ^*p×p*^ is the identity matrix. In banded ridge regression, features are grouped into different feature spaces of size *p*_*i*_, and a different regularization hyperparameter *λ*_*i*_ > 0 is used for each feature space *i*. The weight vector is then computed with *b*^*\**^ = (*X*^⊤^*X* + *D*_*λ*_)^−1^*X*^⊤^*y*, where *D*_*λ*_ ∈ ℝ^*p×p*^ is a diagonal matrix with *λ*_*i*_ repeated *p*_*i*_ times. In banded ridge regression, the optimal *λ*_*i*_ for each feature space are learned by cross-validation. Banded ridge regression is thus able to select the feature spaces relevant for prediction, and optimally scales them to maximize cross-validated prediction accuracy.

Banded ridge regression has been proposed multiple times in the scientific literature, either with a Bayesian approach [Perrakis et al., 2020; Ignatiadis and Lolas, 2020; van Nee et al., 2021] or with a frequentist approach [Nunez-Elizalde et al., 2019; van de Wiel et al., 2021]. In the Bayesian approach, a prior assumption is made on the regularization hyperparameter distribution to simplify the optimization problem. In the frequentist approach, no prior assumption is made on the hyperparameter distribution, and the hyperparameters are optimized based on the average cross-validated prediction accuracy. The present paper uses the frequentist approach. Note that banded ridge regression is sometimes called *group-regularized ridge regression* [van de Wiel et al., 2021]. However, the present paper uses the term *banded ridge regression* [Nunez-Elizalde et al., 2019] because in Neuroscience studies, “group” is most often used for groups of subjects and not groups of features.

### 2.2 Banded ridge regression - model definition

In banded ridge regression, the features are grouped into *m* feature spaces. A feature space *i* is formed by a matrix of features 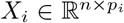, with *n* samples and *p*_*i*_ features. Each feature space is associated with a different regularization hyperparameter *λ*_*i*_ > 0. To model brain activity *y* ∈ ℝ^*n*^ on a particular voxel, banded ridge regression computes the weights 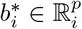 (concatenated into *b*^*\**^ ∈ ℝ^*p*^ with *p* = Σ_*i*_ *p*_*i*_) defined as

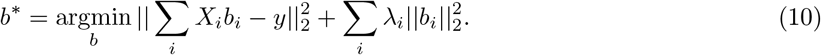

Similarly to ridge regression, the parameters 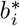 are learned on the training data set, and the hyperparameters *λ*_*i*_ are learned by cross-validation [Nunez-Elizalde et al., 2019]. Because banded ridge regression has multiple hyperparameters *λ*_*i*_, the optimization is more challenging than in ridge regression. (Different strategies for optimizing banded ridge regression are presented in Section 3.)

The benefit of banded ridge regression is its ability to optimize separately the regularization strength of each feature space. This optimization is equivalent to optimizing the relative scale of each feature space. To see this equivalence, the model definition can be rewritten with a change of variable 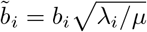 and 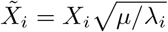 where *µ* > 0 is an arbitrary scalar. The equation becomes 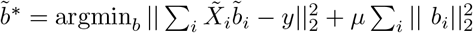. One can then rewrite 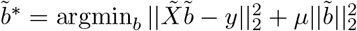 by concatenating 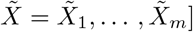 and 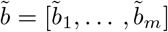. This equation reveals that banded ridge regression is equivalent to performing ridge regression on the scaled features 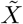 [Hansen, 1998; Nunez-Elizalde et al., 2019], with a regularization strength *µ*.

### 2.3 A feature-space selection mechanism

Banded ridge regression extends ridge regression to adapt the regularization hyperparameter for each feature space. Importantly, the optimization of multiple regularization hyperparameters also leads to a feature-space selection mechanism. Feature-space selection consists in selecting a subset of feature spaces to be used in the regression, and ignoring the rest of the feature spaces. When a feature space has little predictive power, removing it from the joint model can improve generalization performance by reducing the possibility of overfitting to these features. During cross-validation, banded ridge regression is able to learn to ignore some feature spaces, in order to improve generalization performance. To ignore a feature space *i*, banded ridge regression sets a large value for the regularization hyperparameter *λ*_*i*_, and the coefficients 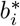 are shrunk toward zero. This process effectively removes the feature space *i* from the model.

With its feature-space selection mechanism, banded ridge regression can select feature spaces that have good predictive power, and suppress those that have little predictive power or that are redundant with other feature spaces. For example, given three correlated feature spaces, banded ridge regression is able to automatically choose between (a) splitting the variance between all three feature spaces, (b) ignoring a redundant feature space and only using the two other ones, or (c) using only one feature space. This example is demonstrated in simulated examples in Appendix 2.4. The feature-space selection thus follows the principle of parsimony (Occam’s razor), while allowing multiple feature spaces to be used jointly.

In voxelwise encoding models, the feature-space selection can be performed independently on each voxel. The joint model is thus able to select different feature spaces depending on the voxel. To illustrate this powerful property, [Nunez-Elizalde et al., 2019] presented the following example (although not explaining it as a feature-space selection mechanism). In their analysis, two feature spaces were used to predict brain activity during a visual task recorded with fMRI. The first feature space was a low-level representation of motion in the stimulus (motion energy, [Nishimoto et al., 2011]), and the second feature space was a semantic representation of the objects and actions present in the scene [Huth et al., 2012]. Because the dataset used naturalistic stimuli, the two feature spaces are correlated. Because of these correlations, a regression model fit with the semantic feature space had significant prediction accuracy in low-level visual areas. This model could lead to the incorrect interpretation that low-level visual areas contain visual semantic representations. To address this issue, both feature spaces were used in a joint model fit with banded ridge regression. The joint model ignored the semantic feature space in low-level visual areas, ignored the low-level feature space in visual semantic areas, and used both feature spaces in other visual areas. This example illustrates how having a different feature-space selection in each voxel can avoid incorrect interpretations due to correlations in naturalistic stimuli.

### 2.4 Example A: Feature-space selection in simulated examples

To illustrate the feature-space selection mechanism of banded ridge regression, four simulated signals were generated from three simulated feature spaces. In two simulated signals, only a subset of the three feature spaces had predictive power. In two other simulated signals, some feature spaces were redundant with other feature spaces. The goal of these simulations is to demonstrate that banded ridge regression is able to ignore the non-predictive or redundant feature spaces.

#### Compared models

Three different regression models are compared. For the first model, a separate ridge regression model was fit on each feature space independently, and the best feature space per voxel was selected based on the cross-validation prediction accuracy. This first model implements the simple winner-take-all model comparison procedure described in Section 1.4, and it does not use a joint model or variance decomposition. The second model is a ridge regression model fit jointly on all feature spaces. This second model takes into account a possible complementarity of feature spaces, but the ridge regularization does not lead to any feature-space selection. The third model is a banded ridge regression model fit on all feature spaces jointly. This third model takes into account a possible complementarity of feature spaces, and the banded ridge regularization leads to a feature-space selection as described in Section 2.3.

For each model, prediction accuracy was measured by the explained variance (*R*^2^-score) on a separate test set. The explained variance was then decomposed over the three feature spaces using the product measure. The results are shown in Figure 3.

**Figure 3:**
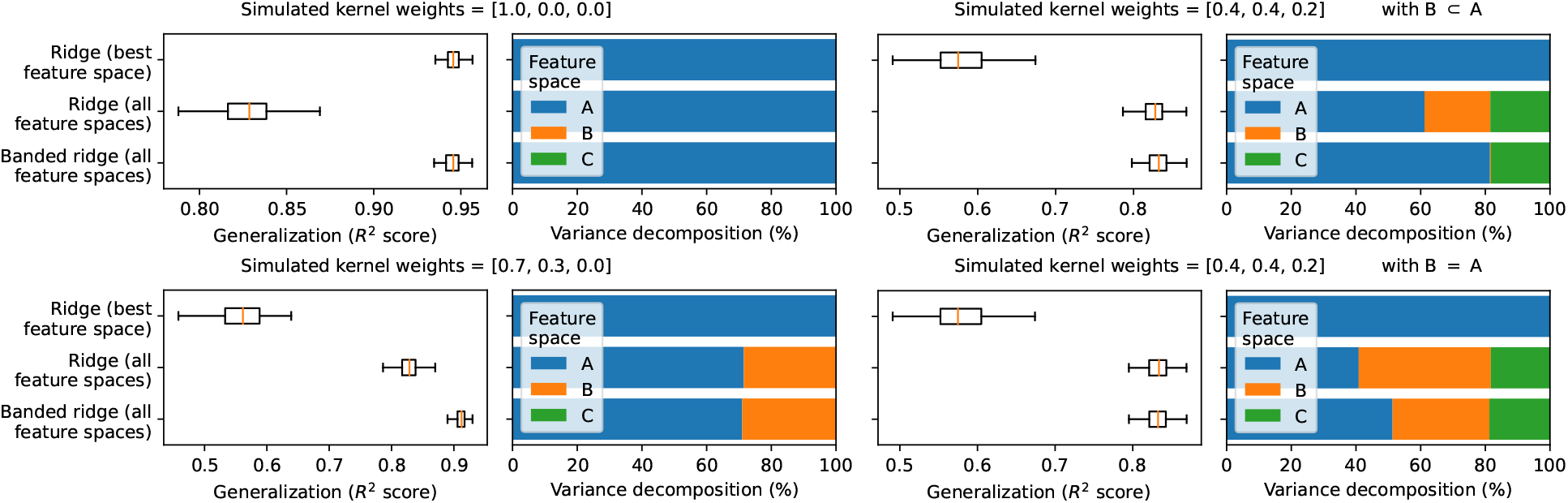
Feature-space selection in four simulated examples. To demonstrate the feature-space selection of banded ridge regression, three different models were fit on four simulated examples, each containing three feature spaces (A, B, C). For each model, prediction accuracy was measured by the explained variance (*R*^2^-score) on a separate test set, using a box plot to show the distribution of score over 100 repetitions of the simulation. The box plot indicates the distribution median (orange bar), the first and third quartiles (box borders), and the 5-th and 95-th percentiles (whiskers). The explained variance was decomposed over the three feature spaces using the product measure, using a geometric median to find a typical decomposition over 100 repetitions. **(Top left)** In the first simulation, the only predictive feature space is A. All three models recover the correct variance decomposition. The prediction accuracy is lower for the ridge regression model fit on all feature spaces than for the two other models, because the model is affected by noise in the non-predictive feature spaces (B, C). **(Bottom left)** In the second simulation, two feature spaces have predictive power (A, B). The ridge regression model fit on the best feature space only uses feature space A, which leads to low prediction accuracy. The ridge regression model fit on all feature spaces recover the correct variance decomposition, but its prediction accuracy is affected by noise in the non-predictive feature space C. The banded ridge regression model fit on all feature spaces recovers the correct variance decomposition, and correctly ignores the non-predictive feature space C to maximize prediction accuracy. **(Top right)** In the third simulation, all three feature spaces are predictive, but feature space B is a subset of A. The ridge regression model fit on all feature spaces uses all three feature spaces, whereas the banded ridge regression model fit on all feature spaces only uses feature spaces A and C, detecting that B is redundant with A. **(Bottom right)** In the fourth simulation, all three feature spaces are predictive, but feature spaces A and B are identical. The ridge regression model fit on all feature spaces uses A and B equally. The banded ridge regression model fit on all feature spaces uses either (A, B, C), or only (A, C), or only (B, C), depending on the repetition, because all three possibilities lead to similar cross-validation scores.

#### Simulations with non-predictive feature spaces

In the first two simulations, some feature spaces are non-predictive. The ridge regression model fit on the best feature space is not affected by these non-predictive feature spaces, but it cannot use multiple feature spaces jointly. The ridge regression model fit on all feature spaces is able to use multiple feature spaces jointly, but it tends to overfit the non-predictive feature spaces, which reduces its prediction accuracy. The banded ridge regression model fit on all feature spaces leads to the best prediction accuracy, because it is able to use multiple feature spaces jointly while ignoring the non-predictive ones.

#### Simulations with non-predictive feature spaces

In the other two simulations, some feature spaces are redundant with other feature spaces. The ridge regression model fit on all feature spaces is not affected by this redundancy, and the shared variance is equally decomposed between the feature spaces. In the case of the banded ridge regression model fit on all feature spaces, if one feature space is strictly better than another, it is selected and the other is ignored. However, if two feature spaces are identical, banded ridge regression is not able to choose between using (i) the first feature space, (ii) the second feature space, or (iii) both at the same time. All three possibilities lead to similar cross-validation scores. The final variance decomposition might thus depend on minor elements such as the initialization in hyperparameter gradient descent, the Dirichlet concentration in the hyperparameter random search, or small variations between the two feature spaces.

### 2.5 Quantifying feature-space selection

In Section 2.6 and Section 2.7, two additional examples are proposed to demonstrate the feature-space selection mechanism in banded ridge regression. For the sake of demonstration, these examples need a metric to quantify the number of feature spaces effectively used in the model. One candidate metric is to count the number of feature spaces with a non-zero effect on the prediction. However, because the hyperparameters optimized by banded ridge regression never reach infinity, the coefficients *b*^*\**^ are never exactly zero, even for feature spaces that are effectively ignored. For this reason, all feature spaces have non-zero values in the variance decomposition, and one cannot simply count the number of non-zero values.

To address this issue, we leverage a metric called the *effective rank* [Roy and Vetterli, 2007; Langeberg et al., 2019; Bartlett et al., 2020]. Starting from a variance decomposition *ρ* ∈ ℝ^*m*^ (for example, using the product measure as described in Section 1.6), negative values are first clipped to zero, and the variance decomposition vector is normalized to sum to one. Then, the effective rank is computed, as defined in Roy and Vetterli [2007] (see Appendix A.1 for alternative definitions)

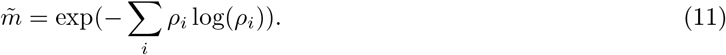

The effective rank is a continuous function of the *m* variance decomposition values *ρ*_*i*_, with values in the interval [1, *m*]. The effective rank is equal to *k* when the variance decomposition is equally split between *k* feature spaces. Thus, the effective rank is used in the following two examples to quantify the feature-space selection in banded ridge regression. (Note that the effective rank is independent of the number of features in each feature space. The following two examples only use the effective rank to measure *feature-space* selection.)

### 2.6 Example B: Feature-space selection over 22 feature spaces

This subsection presents a second example to demonstrate the feature-space selection mechanism in banded ridge regression. The example uses an fMRI dataset recorded while a subject watched several short films [Nunez-Elizalde et al., 2018]. The dataset contains brain responses of 85483 cortical voxels recorded every two seconds. The brain responses were modeled using 22 feature spaces extracted from the stimuli [Nunez-Elizalde et al., 2018]. Examples of feature spaces are the spatio-temporal wavelet transform magnitudes of visual stimuli [Nishimoto et al., 2011], hierarchical classification of objects present in the scenes [Huth et al., 2012], and word embeddings of speech content [Huth et al., 2016] (see Appendix A.2 for a list of all 22 feature spaces). Of the 4379 total time samples, 3572 samples were used in the train set and 807 samples in the test set.

#### Compared models

This example compares the same three models used in Example A (Section 2.4): (1) ridge regression fit on the best predicting feature space per voxel, (2) ridge regression fit jointly on all feature spaces, and (3) banded ridge regression fit jointly on all feature spaces.

#### Hyperparameter optimization

For the ridge regression models, the regularization hyperparameter *λ* was optimized over a grid-search with 30 values spaced logarithmically from 10^−5^ to 10^20^. This range was large enough to ensure that the algorithm explored many different possible regularization strengths for each feature space. For banded ridge regression, the hyperparameters were optimized with the hyperparameter random-search solver described in Section 3.5, using the settings described in Section 3.8.

Because neighboring time samples are correlated in fMRI recordings, any cross-validation scheme should avoid separating correlated time samples into the train and validation sets. Therefore, time samples recorded consecutively were grouped together into “runs”, and a leave-one-run-out cross-validation scheme was used to optimize the hyperparameters. After hyperparameter optimization, all models were refit on the entire training dataset with the optimized hyperparameters.

#### Prediction accuracy comparison

The three models were first compared in terms of prediction accuracy. To estimate prediction accuracy, the explained variance (*R*^2^) was computed on a separate test set that was not used during model fitting. The results are shown in Figure 4. The left panel compares prediction accuracy of the banded ridge regression model to the winner-take-all ridge regression model. The banded ridge regression outperforms the winner-take-all model, especially for voxels that have relatively more accurate predictions. The right panel compares prediction accuracy of the banded ridge regression model to the ridge regression model that learned a single regularization value for all 22 feature spaces. Here again, the banded ridge regression model outperforms the ridge regression model. These results show that prediction accuracy is improved when multiple feature spaces are fit simultaneously and when each space is fit using an optimal regularization parameter for that specific space.

**Figure 4:**
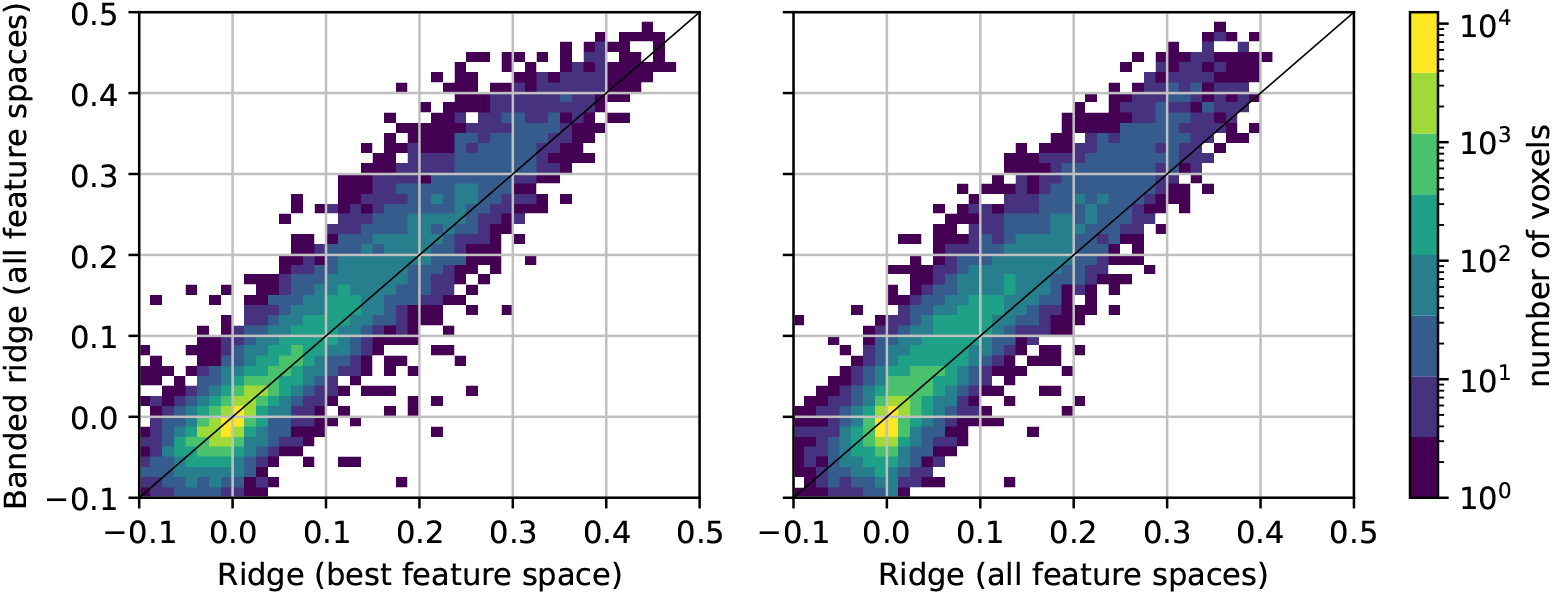
Comparison of prediction accuracy of three different regression models. Three different models were fit using 22 feature spaces to predict brain activity of a subject watching several short films [Nunez-Elizalde et al., 2018]. The prediction accuracy of each model was measured by the explained variance (*R*^2^ score) computed on a separate test set. Each panel compares two models using a 2D histogram over voxels. The diagonal indicates equal performance for both models. **(Left)** Comparison of ridge regression fit on the best feature space per voxel (horizontal) to banded ridge regression fit on all feature spaces (vertical). The mass of the histogram is above the diagonal, indicating better performance for banded ridge regression. It shows that prediction accuracy was improved when multiple feature spaces were fit simultaneously with a separate hyperparameter per feature space, instead of using only the best predicting feature space per voxel. **(Right)** Comparison of ridge regression fit on all feature spaces (horizontal) to banded ridge regression fit on all feature spaces (vertical). The mass of the histogram is again above the diagonal, indicating better performance for banded ridge regression. In both cases, learning a separate regularization per feature space improves the performance of the regression model, especially for voxels that have relatively more accurate predictions.

#### Effective rank comparison

We propose that banded ridge regression produces more accurate predictions than ridge regression because banded ridge regression performs feature-space selection. To quantify feature-space selection, the effective rank (see Section 2.5) was computed on each voxel with positive explained variance. The effective rank gives a continuous measure of the number of feature spaces used in the model.

The top row of Figure 5 compares the effective rank 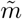 (horizontal) and the explained variance *R*^2^ (vertical) on each of the three models. In the winner-take-all ridge regression model, the effective rank is always equal to 1, because the model only uses the best feature space per voxel. In the ridge regression model fit on all 22 feature spaces, most values of the effective rank are between 2.4 and 10.3 (5th and 95th percentiles) for the best-predicted voxels (*R*^2^ > 0.05). These effective rank values are high because ridge regression does not remove redundant feature spaces, so many feature spaces contribute to the explained variance. In the banded ridge regression model, most values of the effective rank are between 1.0 and 3.7 (5th and 95th percentiles) for the best-predicted voxels (*R*^2^ > 0.05). These results indicate that banded ridge regression effectively uses fewer feature spaces than ridge regression, but that banded ridge regression can still use multiple feature spaces jointly when necessary.

**Figure 5:**
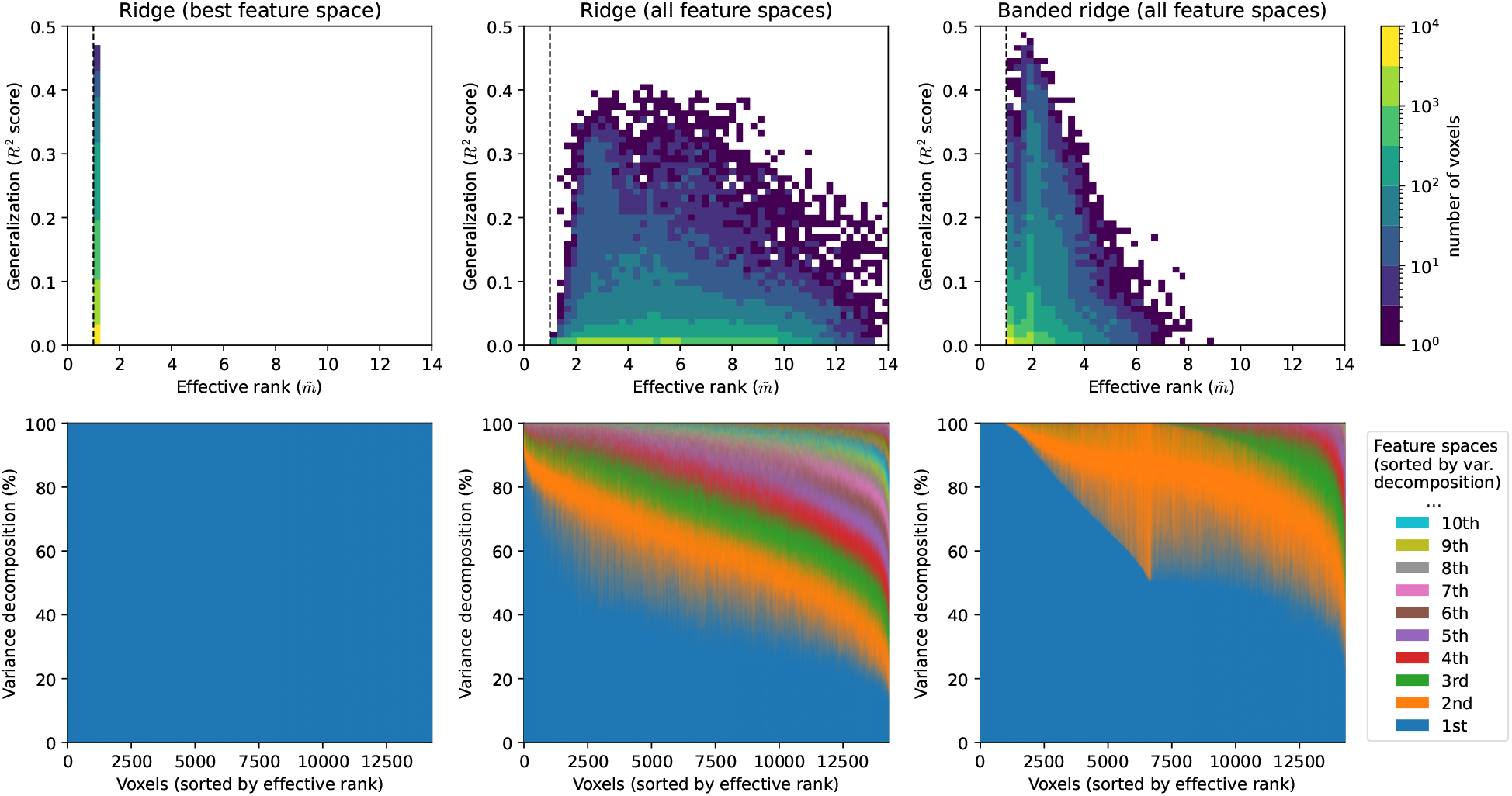
Feature-space selection over 22 feature spaces in three different regression models. To describe the feature-space selection mechanism in banded ridge regression, feature-space selection is quantified using the effective rank 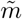 The effective rank is a measure that quantifies the number of feature spaces effectively used by a model in a particular voxel. The three columns correspond to three different models. The (top row) shows a 2D histogram over voxels with *R*^2^ ≥ 0, comparing the effective rank 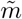 (horizontal) and the explained variance *R*^2^ (vertical). This plot describes the range of values obtained with the effective rank. The (bottom row) shows the variance decomposition over feature spaces, for voxels with *R*^2^ ≥ 0.05. For each voxel, the 22 feature spaces are sorted by contribution to the variance decomposition. Then, voxels are sorted by effective rank 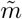 The variance decomposition of each voxel is then displayed using stacked areas. This plot gives an intuitive view of what is captured by the effective rank metric. **(Left)** Ridge regression on the best feature space. On each voxel, the best performing feature space is selected. The effective rank is thus always equal to 1. This winner-take-all selection ignores the possibility that different feature spaces might be complementary on some voxels. **(Middle)** Ridge regression. Many feature spaces contribute to part of the variance, as captured by larger effective rank values. This model does not remove redundant feature spaces. **(Right)** Banded ridge regression. This model contains an implicit feature-space selection mechanism, as indicated by the lower effective rank values. This feature-space selection helps remove redundant feature spaces, while still allowing multiple feature spaces to be used jointly.

#### Variance decomposition comparison

In the previous paragraph, the feature-space selection of banded ridge regression is measured with the effective rank. To understand more intuitively what the effective rank measures, the variance decomposition results can be visualized for voxels of increasing effective rank. The variance decomposition results were computed by first selecting a subset of best-predicted voxels (*R*^2^ > 0.05). Then, the product measure (as defined in Section 1.6) was used to compute the variance decomposition, *ρ* ∈ ℝ^*m*^, of these voxels. In this setting, negative values, if present, were very small, and could be clipped to 0. The results were normalized to sum to one to obtain the relative contribution of each feature space to the explained variance. Next, for each voxel separately, the 22 feature spaces were sorted by their relative contribution (*ρ*_1_ ≥ *ρ*_2_ ≥ … > *ρ*_*m*_).

The bottom row of Figure 5 shows the normalized variance decomposition as stacked areas, where voxels are sorted along the horizontal axis by their effective rank 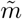 For example, the first stacked area (dark blue) corresponds to the relative contribution of the first feature space (*i*.*e*. the feature space with the highest relative contribution). In the winner-take-all ridge regression model, the first feature space contributes 100% of the explained variance in every voxel. In the ridge regression model fit on all 22 feature spaces, the first feature space contributes between 20% and 90% of the explained variance. In banded ridge regression, the first feature space contributes between 30% and 100% of the explained variance. Overall, these results indicate that banded ridge regression leads to fewer feature spaces being effectively used than with ridge regression, but that banded ridge regression can still use multiple feature spaces jointly when necessary.

### 2.7 Example C: Feature-space selection over 7 neural network layers

This subsection presents a third example to further demonstrate the feature-space selection mechanism in banded ridge regression. It reuses part of the results described in a previous conference publication of our lab [Dupré la Tour et al., 2021]. This example uses an fMRI dataset recorded while a subject watched short movie clips [Nishimoto et al., 2011]. The dataset contains brain responses of 73211 cortical voxels recorded every two seconds. The brain responses were modeled using 7 feature spaces extracted from the stimuli using intermediate layers of a pre-trained convolutional neural network (CNN) (see the feature extraction paragraph below). Of the 3870 total time samples, 3600 samples were used in the train set and 270 samples in the test set.

Several previous studies have used these sorts of features to compare visual representations in CNNs versus the visual cortex. For example, several studies have used this approach to argue that early CNN layers best predict brain activity in low-level visual areas; late layers best predict brain activity in intermediate and higher-level visual areas; and that the CNN layer that best describe visual tuning changes gradually across the cortical visual hierarchy [Yamins et al., 2014; Agrawal et al., 2014; Cichy et al., 2016; Güçlü and van Gerven, 2015; Eickenberg et al., 2016; Yamins and DiCarlo, 2016; Wen et al., 2018; St-Yves and Naselaris, 2018; Zhang et al., 2019; Zhuang et al., 2021; Nonaka et al., 2021]. This approach has also been used to map the hierarchical representation of features in speech [Kell et al., 2018; Millet and King, 2021] and language [Jain and Huth, 2018; Toneva and Wehbe, 2019].

One limitation of using CNNs as sources of features for modeling brain data is that the features represented in different layers of a CNN are usually strongly correlated. Thus, different encoding models trained with features from different CNN layers have similar prediction accuracies [Eickenberg et al., 2016; Jain and Huth, 2018; Toneva and Wehbe, 2019; Wang et al., 2019]. It is then not clear how to interpret these different models, and disentangle their relative contributions. Most studies ignore this issue and select the best-predicting layer for each voxel [Agrawal et al., 2014; Güçlü and van Gerven, 2015; Eickenberg et al., 2016; Wen et al., 2018; Kell et al., 2018; Schrimpf et al., 2020; Zhang et al., 2019], but this winner-take-all approach is oversimplistic, not robust to noise, and ignores potential complementarities between layers. Some studies use variance partitioning [Groen et al., 2018] or canonical component analysis [Yang et al., 2019] to disentangle the different layers, but these approaches cannot disentangle more than two or three layers. The present example demonstrates how variance decomposition and banded ridge regression can be used to disentangle the contributions of many different CNN layers. The obtained contributions of all CNN layers can then be aggregated into a continuous layer mapping with increased smoothness over the cortical surface.

#### Feature extraction

To extract features, each image of the stimuli was first presented to a pretrained image-based CNN “Alexnet” [Krizhevsky et al., 2012]. Then, the activations of a CNN layer were extracted (after ReLU and max-pooling layers). Features were extracted frame by frame, and thus needed to be down-sampled to the brain imaging sampling frequency (typically 0.5 Hz). To avoid removing high-frequency information, the layer activations were filtered with eight complex-valued band-pass filters (of frequency bands [0, 0.5], [0.5, 1.5], …, [6.5, 7.5] Hz) (see more details in [Dupré la Tour et al., 2021]). These band-pass filters were designed to preserve information from different temporal frequency bands, similarly to spatio-temporal features from [Nishimoto et al., 2011]. Then, the amplitude of each filtered complex-valued signal was computed to extract the envelope of the signal (the envelope is the slowly varying amplitude modulation of the high-frequency signal). Next, the envelope was down-sampled to the fMRI sampling frequency with an anti-aliasing low-pass filter. Next, a compressive nonlinearity *x* ↦ log(1 + *x*) was applied, and features were centered individually along the train set. Finally, to account for the delay between the stimulus and the hemodynamic response, features were duplicated with four temporal delays of *d* ∈ {2, 4, 6, 8} seconds. This process was repeated on seven CNN layers, to create seven separate feature spaces.

#### Compared models

This example compares the same three models used in Example A (Section 2.4): (1) ridge regression fit on the best predicting layer per voxel, (2) ridge regression fit jointly on all layers, and (3) banded ridge regression fit jointly on all layers.

#### Hyperparameter optimization

The hyperparameter optimization settings were identical to the ones in Example B (Section 2.6).

#### Prediction accuracy comparison

The three models were first compared in terms of prediction accuracy. To estimate prediction accuracy, the explained variance (*R*^2^) was computed on a separate test set that was not used during model fitting. A detailed comparison of the prediction accuracy of these three models can be found in [Dupré la Tour et al., 2021] and is omitted here for the sake of brevity. In particular, the study shows that the banded ridge regression model outperforms both the ridge regression model fit on the best feature space, and the ridge regression model fit on all the feature spaces.

#### Feature-space selection comparison

Again, we propose that banded ridge regression produces more accurate predictions than ridge regression because banded ridge regression performs a feature-space selection. To visualize feature-space selection, the explained variance was decomposed over the seven feature spaces (*i*.*e*. over the seven CNN layers) using the product measure (described in Section 1.6).

Figure 6a shows examples of this variance decomposition on four well-predicted voxels. In ridge regression fit on the best layer, because the model uses only one layer, the variance decomposition is concentrated in a single contribution, and ignores potential layer complementarity. In ridge regression fit jointly on all layers, the variance decomposition is spread across all seven layers, showing that the model uses all layers to make the predictions. In banded ridge regression, the variance decomposition is concentrated on a small number of layers. The model contains a feature-space selection mechanism, leading to sparsity at the layer level, yet allowing multiple layers to be used for complementarity and robustness.

**Figure 6:**
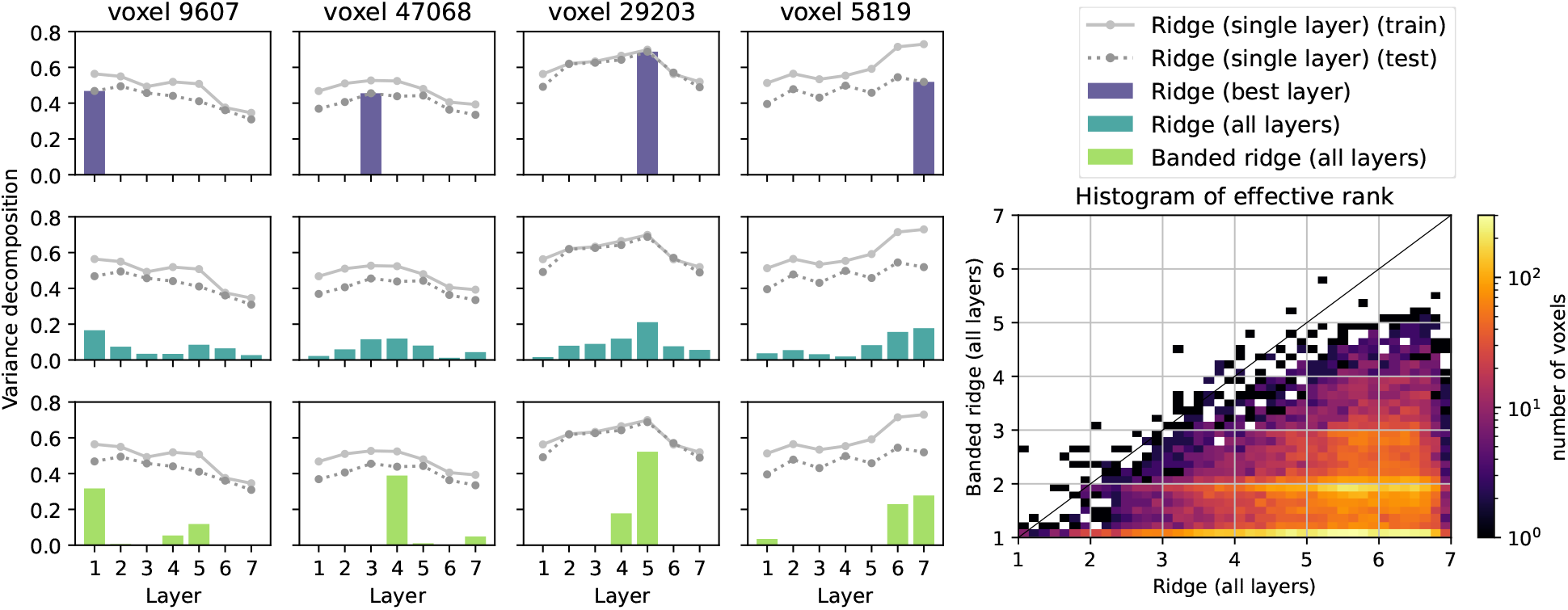
Feature-space selection on seven convolutional neural networks layers. Seven feature spaces were extracted from a movie stimulus based on activations from seven layers of a pretrained CNN (Alexnet [Krizhevsky et al., 2012]). These feature spaces were then used in three different encoding models to predict brain responses to the movie stimulus. The product measure was then used to decompose the explained variance (*R*^2^ score) over the seven layers. **(a)** Comparison of three models in four voxels of brain activity. The first model (*top row*) uses ridge regression fit on the best layer only, ignoring potential complementarity between layers. The second model (*middle row*) uses ridge regression fit jointly on all layers. The variance decomposition shows that this second model uses almost all layers to make the predictions. The third model (*bottom row*) uses banded ridge regression fit jointly on all layers. The variance decomposition shows that this third model uses only two or three layers to make the predictions. Indeed, banded ridge regression performs a feature-space selection, yet allows multiple layers to be used simultaneously. Two gray lines are also given as references, showing the prediction accuracy of a ridge regression model fit on a single layer. These two prediction accuracies are computed either on the train set or on the test set. **(b)** To quantify the number of layers effectively used by each model, the effective rank was computed on the variance decomposition. The 2D histogram of effective rank over voxels shows that banded ridge regression (*vertical*) used fewer feature-spaces (smaller effective rank) than ridge regression (*horizontal*) in almost all voxels.

To quantify feature-space selection, the effective rank (see Section 2.5) was computed over all significantly predicted voxels (*p* < 0.01, permutation test). Comparing the distribution of effective rank over voxels, Figure 6b shows that on each voxel, banded ridge regression used fewer layers than ridge regression.

#### Layer mapping comparison

The three models can also be compared in terms of layer mapping. To describe similarities of representations between CNNs and the visual cortex, it is common to use the index of the best predicting layer as a layer mapping metric on each voxel [Yamins et al., 2014; Agrawal et al., 2014; Cichy et al., 2016; Güçlü and van Gerven, 2015; Eickenberg et al., 2016; Yamins and DiCarlo, 2016; Wen et al., 2018; Dupré la Tour et al., 2021]. As many studies have shown, plotting this layer mapping on the cortex reveals a gradient over the cortical surface (Figure 7a). However, because of correlation between layers, two layers can lead to very similar prediction accuracies. In this case, the best-layer selection picks one layer over the other, even if the prediction accuracy difference is small. The best-layer selection is thus non-robust.

**Figure 7:**
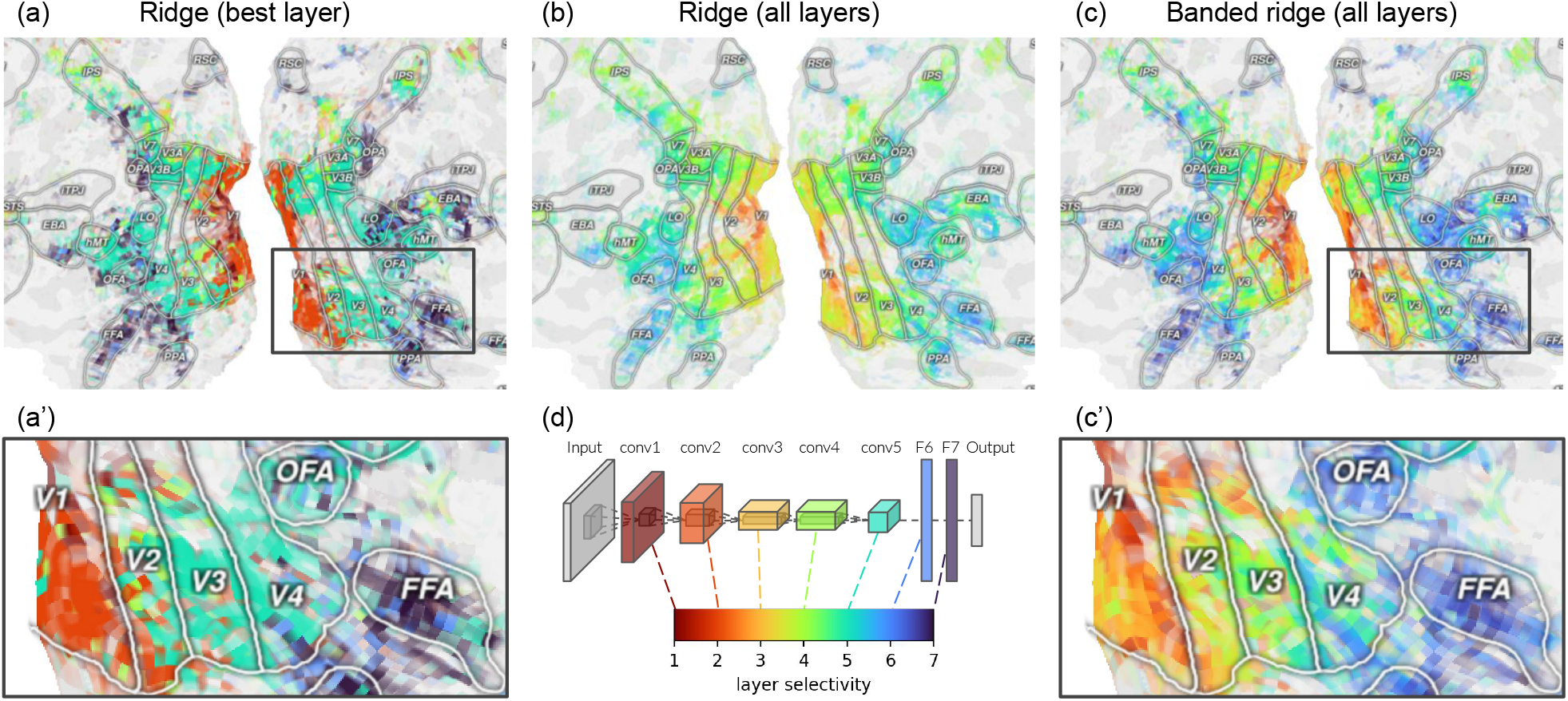
Mapping convolutional neural network layers to the visual cortex. Seven feature spaces were extracted from a movie stimulus based on activations from seven layers of a pretrained CNN (Alexnet [Krizhevsky et al., 2012]). These feature spaces were then used in three different encoding models to predict brain responses to the movie stimulus. Using the fit models, these layers were then mapped to each voxel in the visual cortex. The layer mapping of each voxel is computed as a weighted average of the layer indices, weighted by the decomposition over layers of the explained variance. For visualization, the layer mapping is projected on a flattened cortical surface using Pycortex [Gao et al., 2015]. **(a)** The first model uses ridge regression fit on the best layer only. This winner-take-all approach gives a non-robust estimate of layer mapping, because the best-layer selection can flip from one layer to another due to small variations in prediction accuracy. **(b)** The second model uses ridge regression fit jointly on all layers. It leads to a continuous measure of layer mapping, with a smooth gradient over the cortical surface. However, fitting the joint model with ridge regression gives a biased estimate of layer mapping toward middle values, because its variance decomposition tends to use all layers. **(c)** The third model uses banded ridge regression fit jointly on all layers. Fitting with banded ridge regression mitigates the bias toward middle values thanks to its feature-space selection. **(a’, c’)** Zoomed views of (a, c). **(d)** Correspondence with the CNN architecture. The layer mapping is derived from the layer indices of a pretrained Alexnet model.

To solve this issue, we extend the definition of layer mapping using a weighted average of layer indices, where the weights are the variance decomposition results. Starting from a variance decomposition *ρ* ∈ ℝ^*m*^, where *m* is the number of layers, negative values are clipped to zero, and *ρ* is normalized to sum to one. Then, the layer mapping is computed as a weighted average 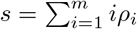 This weighted average leads to a *continuous* layer mapping, which increases both its robustness and its ability to describe gradual changes over the cortical surface.

The weighted average can be computed on any joint model, either fit with ridge regression (Figure 7b) or with banded ridge regression (Figure 7c). In both cases, the continuous layer mapping results in a smoother gradient of layer mapping across the cortical surface. However, the layer mapping estimated by the ridge regression model is biased toward middle values, because the ridge regression model tends to use all layers. Banded ridge regression reduces this bias toward middle values thanks to its feature-space selection mechanism.

### 2.8 Other models with feature-space selection mechanisms

Feature space selection is an important aspect of many regression algorithms. In the statistics literature, feature-space selection is called *group sparsity*. This subsection reviews a number of models that induce group sparsity, and it describes how each model is related to banded ridge regression.

#### Automatic relevance determination

Automatic relevance determination [MacKay, 1994; Neal, 1995] is a framework initially developed for Bayesian neural networks [MacKay, 1992b], a class of neural networks where all parameters have explicit prior distributions. In automatic relevance determination, a separate hyperparameter is optimized on each input feature (or each group of features). This hyperparameter optimization induces a feature (or feature-space) selection, automatically determining which features (or feature spaces) are most relevant.

Automatic relevance determination was later applied to Bayesian ridge regression [Box and Tiao, 1973; MacKay, 1992a], a formulation of ridge regression where all parameters and hyperparameters have explicit prior distributions. Using automatic relevance determination on Bayesian ridge regression produces a model called sparse Bayesian learning [Tipping, 2001; Wipf and Nagarajan, 2007]. Sparse Bayesian learning is the direct Bayesian equivalent of banded ridge regression. The main benefit of the Bayesian framework is that all hyperparameters have explicit priors, which leads to efficient optimization solvers without cross-validation [van Nee et al., 2021; Perrakis et al., 2020; Ignatiadis and Lolas, 2020]. However, the lack of cross-validation increases the risk of overfitting. In contrast, banded ridge regression does not make assumptions about hyperparameter prior distributions. To optimize the hyperparameters, banded ridge regression uses cross-validation to select hyperparameters with the best generalization performance.

#### Multiple-kernel learning

Multiple-kernel learning [Lanckriet et al., 2004; Bach et al., 2004] is a framework that was developed for use with support vector machines [Boser et al., 1992], a class of versatile models for classification and regression. In support vector machines, the features *X* are only used through a pairwise similarity matrix *K*(*X*) called the kernel matrix. In the kernel matrix, each entry is a pairwise similarity between two samples *K*(*X*)_*ij*_ = *k*(*X*_*i*_, *X*_*j*_), where *k* is a kernel function. Different kernel functions can be used depending on the problem at hand, for example the linear kernel *k*(*x, y*) = *x*^⊤^*y*, or the radial-basis-function kernel *k*(*x, y*) = exp(−||*x* − *y*||^2^*/*2*σ*^2^) of size *σ*. Because different kernels (or different kernel hyperparameters) can lead to different predictive performances, a cross-validation is generally performed to select the best kernel. To leverage the strength of multiple kernels jointly, the multiple-kernel learning framework uses a weighted kernel *K* = Σ_*i*_ *γ*_*i*_*K*_*i*_, where *γ*_*i*_ are positive kernel weights learned by the model. If a kernel weight *γ*_*i*_ has a value close to zero, the associated kernel *K*_*i*_ has little effect in the model. Therefore, learning the kernel weights leads to a kernel-selection mechanism, where only a subset of the kernels are effectively used by the model. If the different kernels are built on different feature spaces, the kernel-selection mechanism corresponds to a feature-space selection mechanism.

This framework has also been applied to ridge regression [Hoerl and Kennard, 1970]. To do so, ridge regression is first re-formulated as a kernel ridge regression [Saunders et al., 1998] with a linear kernel (see more details in Section 3.2). Then, the multiple-kernel framework is used to produce a model called multiple-kernel ridge regression [Bach, 2008]. Interestingly, multiple-kernel ridge regression is *almost* equivalent to banded ridge regression. This quasi-equivalence is described in further details in Appendix A.3.

#### Group lasso

The group lasso [Yuan and Lin, 2006] is a generalization of the lasso [Tibshirani, 1996]. The lasso is a linear regression model similar to ridge regression but with a different regularization term. The lasso is defined as 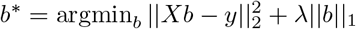, where ||*b*||_1_ = Σ_*j*_ |*b*_*j*_| is the *L*_1_ norm. Due to the geometry of the *L*_1_ norm, the lasso is known to induce a feature selection [Tibshirani, 1996]. The group lasso is a generalization of the lasso which extends this selection mechanism to feature spaces. The group lasso is defined as 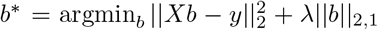 where ||*b*||_2,1_ = Σ_*i*_ ||*b*_*i*_||_2_ is the *L*_2,1_ norm. Due to the geometry of the *L*_2,1_ norm, the group lasso induces a feature-space selection. (To induce both a feature selection and a feature-space selection, the group lasso can be extended to the sparse group lasso [Simon et al., 2013].) A notable variant of the group lasso is the squared group lasso [Bach, 2008], which uses the squared-*L*_2,1_ norm 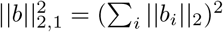. The squared group lasso also induces a feature-space selection.

Interestingly, the squared group lasso has been shown to be equivalent to multiple-kernel ridge regression [Bach, 2008; Rakotomamonjy et al., 2008]. Because banded ridge regression is quasi-equivalent to multiple-kernel ridge regression, it is also quasi-equivalent to the squared group lasso. The details of this quasi-equivalence are described in Appendix A.3.

## 3 Efficient banded ridge regression solvers

As shown in the previous section, banded ridge regression and its feature-space selection mechanism provide a powerful framework for fitting complex encoding models to brain data. However, one obstacle to widespread adoption of this framework is that solving banded ridge regression remains a computational challenge. This section presents several methods for solving banded ridge regression efficiently, and proposes an empirical comparison of these methods. All methods are implemented in an open-source Python package called *Himalaya*.

### 3.1 Ridge regression solver

Before considering banded ridge regression, it is useful to recall how to solve ridge regression. Let *X* ∈ ℝ ^*n×p*^ be the matrix of features, with *n* samples and *p* features, *y* ∈ ℝ ^*n*^ the brain activity vector in one particular voxel, and *λ* > 0 a fixed regularization hyperparameter. Ridge regression [Hoerl and Kennard, 1970] considers the weight vector *b*^*\**^ ∈ ℝ^*p*^ defined by the optimization problem

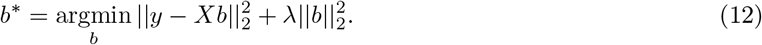

This optimization problem has a closed-form solution *b*^*\**^ = *My*, where *M* = (*X*^⊤^*X* + *λI*_*p*_)^−1^*X*^⊤^ ∈ ℝ^*p×n*^.

#### Efficient solver for multiple voxels

In the closed-form solution above, the resolution matrix *M* does not depend on the brain activity vector *y*. Therefore, when fitting a ridge regression on each voxel independently, the matrix *M* can be precomputed and reused on all voxels. Reusing *M* on all voxels dramatically reduces the computations compared to solving a linear system per voxel. With *t* voxels, the precomputation reduces the complexity of solving the full system from 𝒪 (*p*^3^*t* + *p*^2^*nt*) to 𝒪 (*p*^3^ + *p*^2^*n* + *pnt*). To further decrease the computation time, one can concatenate all voxels into a matrix *Y* and write in matrix form *B*^*\**^ = *MY*. The matrix form uses a matrix-matrix multiplication that is very fast on modern hardware, rather than multiple matrix-vector multiplications that would be inevitably slower. Using these methods, ridge regression can be solved efficiently on large numbers of voxels.

#### Efficient solver for multiple hyperparameters

In ridge regression, the optimal value for the hyperparameter *λ* is unknown, so it is typically selected through a grid-search with cross-validation (see Section 1.2). Thus, there is a need to solve the model efficiently for multiple hyperparameters *λ*. To do so, the training input matrix can be decomposed using the singular value decomposition [Golub and Reinsch, 1971; Hastie and Tibshirani, 2004; Rifkin and Lippert, 2007], *X*_train_ = *USV* ^⊤^, where *U* ∈ ℝ^*n×p*^ is orthonormal, *S* ∈ ℝ^*p×p*^ is di-agonal, and *V* ∈ ℝ^*p×p*^ is orthonormal. (The matrix sizes are given here assuming that *p* ≤ *n*.) Then, for each hyperparameter *λ*, the resolution matrix is given by *M* (*λ*) = *V* (*S*^2^ + *λI*_*p*_)^−1^*SU* ^*T*^. This expression is inexpensive to compute because (*S*^2^ + *λI*_*p*_)^−1^*S* is diagonal. By using the singular value decomposition and looping over *r* values of *λ*, the complexity of computing the resolution matrix decreases from 𝒪 (*p*^3^*r* + *pnr*) to 𝒪 (*p*^2^*nr*). Using this method, ridge regression can be solved efficiently even when the hyperparameter grid search is large. To evaluate each hyperparameter, the predictions on the validation set can be computed with *ŷ*_val_(*λ*) = *X*_val_*M* (*λ*)*Y*_train_ = *X*_val_*V* (*S*^2^ + *λI*_*p*_)^−1^*SU* ^*T*^*Y*_train_. The optimal order to compute this matrix product depends on *p, n*_val_, *n*_train_, and *t*. To further decrease the computation time, the loop over hyperparameters *λ* can also be implemented with a tensor product.

### 3.2 Kernel ridge regression solver

Solving ridge regression as described in Section 3.1 requires 𝒪 (*p*^3^) operations, and this can become expensive when the number of features *p* is large. Fortunately, if the number of features is larger than the number of time samples (*p* > *n*), ridge regression can be reformulated to be solved more efficiently. By the Woodbury matrix identity, *b*^*\**^ can be rewritten *b*^*\**^ = *X*^⊤^(*XX*^⊤^ + *λI*_*n*_)^−1^*y*, or *b*^*\**^ = *X*^⊤^*w*^*\**^ for some *w*^*\**^ ∈ ℝ^*n*^. Given the linear kernel *K* = *XX*^⊤^ ∈ ℝ^*n×n*^, this leads to the equivalent formulation

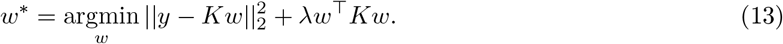

This model is called *kernel ridge regression* [Saunders et al., 1998], and can be used with arbitrary positive semidefinite kernels *K*. When the kernel is linear (*K* = *XX*^⊤^), kernel ridge regression is equivalent to ridge regression.

The kernel formulation has a closed-form solution, *w*^*\**^ = (*K* + *λI*_*n*_)^−1^*y*. Interestingly, this closed-form solution requires inverting a (*n* × *n*) matrix, whereas solving ridge regression requires inverting a (*p* × *p*) matrix. Specifically, the computational complexity of the multiple-voxel case is 𝒪 (*n*^3^ + *pn*^2^ + *n*^2^*t*) for kernel ridge regression, and 𝒪 (*p*^3^ + *p*^2^*n* + *pnt*) for ridge regression. It is thus more efficient to use the kernel formulation when the number of features is larger than the number of time samples *p* > *n*, and it is more efficient to use the regular formulation otherwise.

#### Efficient solver for multiple voxels

Similarly to ridge regression, the resolution matrix *M* = (*K* +*λI*_*n*_)^−1^ can be precomputed and efficiently applied to all voxels. To further decrease the computation time, one can again concatenate all voxels into a matrix *Y* and write in matrix form *w*^*\**^ = *MY*. Using these methods, kernel ridge regression can be solved efficiently on large numbers of voxels.

#### Efficient solver for multiple hyperparameters

Similarly to ridge regression, the hyperparameter *λ* is unknown and needs to be selected with cross-validation. To efficiently solve kernel ridge regression during hyperparameter grid search, *λ*, the training kernel 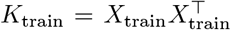 can be diagonalized into *K*_train_ = *UDU* ^⊤^, where *D* ∈ ℝ^*n×n*^ is diagonal, and *U* ∈ R^*n×n*^ is orthogonal. Then, for each hyperparameter candidate *λ*, the resolution matrix can be inverted with (*K* + *λI*_*n*_)^−1^ = *U* (*D* + *λI*_*n*_)^−1^*U* ^⊤^. Here, the kernel diagonalization replaces the matrix inversion with a matrix multiplication, which is less computationally expensive. However, it does not change the computational complexity, which is 𝒪 (*n*^3^*r*) for *r* values of *λ*. Using this method, kernel ridge regression can be solved efficiently even when searching through many different values of the hyperparameters.

Finally, search can also be sped up when computing predictions on the validation set *ŷ*_val_(*λ*) = *K*_val_*U* (*D* + *λI*_*n*_)^−1^*U* ^*T*^*Y*_train_, where 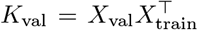 The speed-up depends on the optimal order to compute this matrix product, which depends on *p, n*_val_, *n*_train_, and *t*.

### 3.3 Banded ridge regression solver

As discussed in Section 2.2, banded ridge regression is a natural extension of ridge regression, where the features are grouped into *m* feature spaces. A feature space *i* is formed by a matrix of features 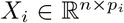, with *n* samples and *p*_*i*_ features, and is associated with a regularization hyperparameter *λ*_*i*_ > 0. To model brain activity *y* ∈ ℝ^*n*^ on a particular voxel, banded ridge regression considers the weights 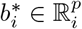 (concatenated into *b*^*\**^ ∈ R^*p*^ with *p* = Σ_*i*_ *p*_*i*_) defined as

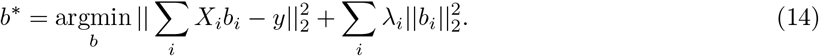

For a fixed hyperparameter vector 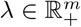 banded ridge regression has a closed-form solution *b*^*\**^ = (*X*^⊤^*X* + *D*_*λ*_)^−1^*X*^⊤^*y*, where *D*_*λ*_ ∈ ℝ^*p×p*^ is a diagonal matrix with *λ*_*i*_ repeated *p*_*i*_ times, and *X* = (*X*_1_, …, *X*_*m*_) ∈ R^*n×p*^ is the concatenation of the *X*_*i*_.

#### Efficient solver for multiple hyperparameters

Because the hyperparameters in banded ridge regression 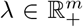 are optimized by cross-validation, hyperparameter optimization must be implemented efficiently. However, the trick used in ridge regression to factorize computations for all hyperparameter candidates cannot be used in banded ridge regression. Indeed, with more than one hyperparameter (*m* > 1), the singular value decomposition of *X* does not codiagonalize *D*_*λ*_. We thus propose two other methods to solve banded ridge regression efficiently: *hyperparameter random search* (described in Section 3.5), and *hyperparameter gradient descent* (described in Section 3.6).

### 3.4 Multiple-kernel ridge regression

For maximal efficiency, the proposed banded ridge regression solvers are described with the kernel formulation of banded ridge regression, called multiple-kernel ridge regression (see Appendix A.3). This formulation uses kernels, and leads to more efficient solvers than banded ridge regression when *p* > *n*. In standard ridge regression, when the number of features is larger than the number of samples *p* > *n*, ridge regression can be solved more efficiently using kernel ridge regression (see Section 3.2). Similarly, when *p* > *n*, banded ridge regression can be solved more efficiently through its equivalent kernel formulation.

As described in Appendix A.3, multiple-kernel ridge regression uses a set of kernels (*K*_1_, …, *K*_*m*_) with *K*_*i*_ ∈ ℝ^*n×n*^, a regularization strength *µ* > 0, and a kernel weight vector *γ* defined on the simplex 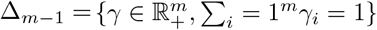. Then, to model brain activity *y* ∈ ℝ^*n*^ on a particular voxel, multiple-kernel ridge regression considers the dual coefficients *w*^*\**^ ∈ ℝ^*n*^ defined by

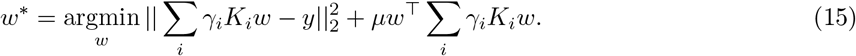

The brain activity is then modeled using *ŷ* = Σ_*i*_ *γ*_*i*_*K*_*i*_*w*^*\**^. To ensure that the kernel formulation is equivalent to banded ridge regression, the kernels need to be defined using one linear kernel 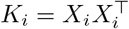 per feature space *i*. The correspondence with banded ridge regression hyperparameters is given by *µ* = (Σ_*i*_1/*λ*_*i*_)^−1^ and *γ*_*i*_ = *µ/λ*_*i*_.

In the following two subsections, two solvers for multiple-kernel ridge regression are proposed. Both methods are summarized in Figure 8, using a simulated toy example. Because the datasets used in this paper are in the setting *p* > *n*, these two methods are described for multiple-kernel ridge regression. To be more efficient when *p* < *n*, similar methods could be derived for banded ridge regression.

**Figure 8:**
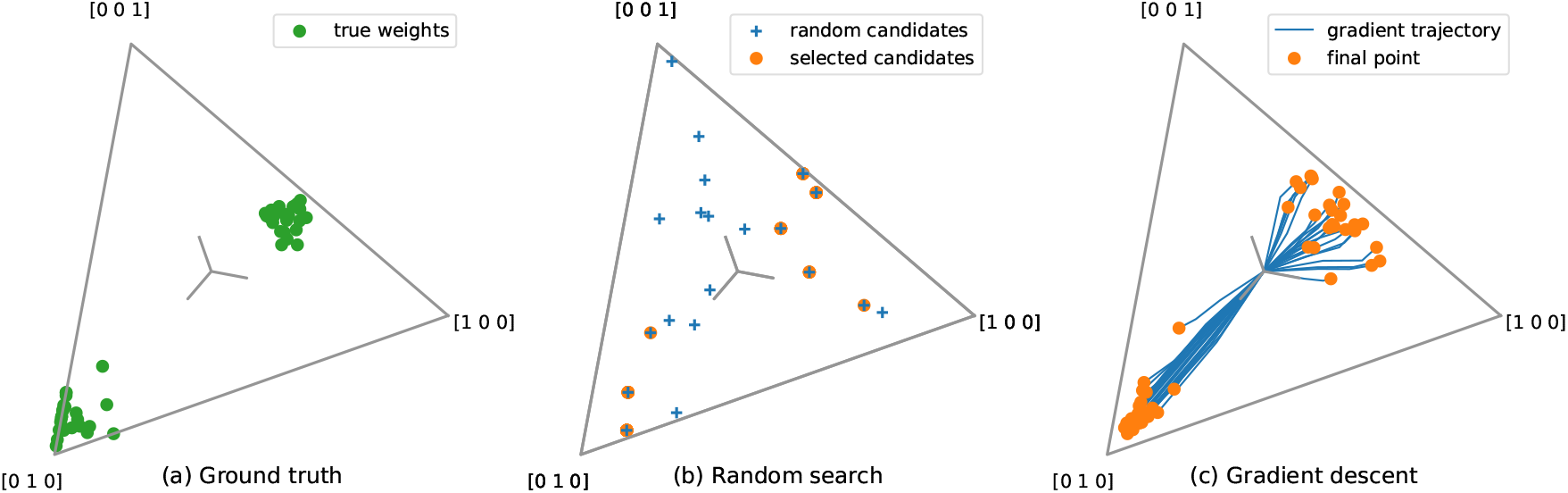
Overview of the two proposed methods to learn hyperparameters in banded ridge regression. In banded ridge regression, a different regularization hyperparameter *λ*_*i*_ > 0 is used on each feature space *i*. All hyperparameters are learned using cross-validation. These hyperparameters can be reparameterized into a regularization strength 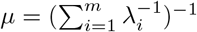 and a set of kernel weights *γ*_*i*_ = *µ/λ*_*i*_. The kernel weight vector is thus defined on the “simplex” 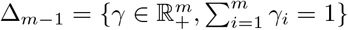. The figure presents the simplex with *m* = 3, which is a 2-dimensional surface in the shape of a triangle. **(a) Ground truth**. Ground-truth kernel weights on the simplex Δ_2_, for 40 different voxels. Brain activity was simulated on 40 voxels, as a function of three feature spaces. On each voxel, the three feature spaces were balanced using a ground-truth kernel-weights vector *γ* ∈ Δ_2_. Note that the ground-truth kernel weights are not necessarily the solution of the optimization problem, because the simulated dataset has a finite size. **(b) Random search**. The random-search method randomly samples candidates from the simplex, and selects for each voxel the candidate leading to the best cross-validation loss. The method is fast because it factorizes computations over all voxels. However, the results directly depend on the random sampling of candidates. **(c) Gradient descent**. The gradient-descent method iteratively optimizes the kernel weights for each voxel independently, using the gradient of the cross-validation loss. The method is more reliable than random search, but it can be slower because it considers each voxel independently.

### 3.5 Method 1: Hyperparameter random search

The first proposed method to solve banded ridge regression is hyperparameter random search [Bergstra and Bengio, 2012]. Hyperparameter random search consists in randomly sampling hyperparameter candidates, and then selecting the candidates that produce the lowest cross-validation error. It is often more efficient than grid search, because the random sampling of hyperparameter space is less redundant. For example, when adding useless hyperparameters to a search, a grid search becomes less efficient, while random search efficiency remains unchanged [Bergstra and Bengio, 2012].

In the specific case of multiple-kernel ridge regression, *m* hyperparameters are optimized, parameterized with *γ* ∈ Δ_*m*−1_ and *µ* > 0. Formally, the training and validation loss functions are defined as

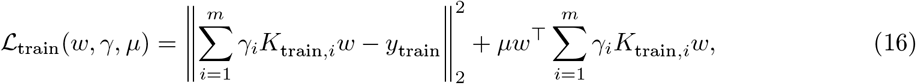

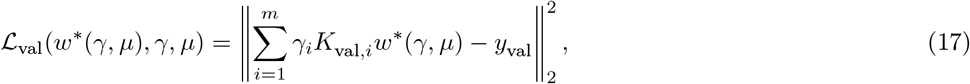

where *w*^*\**^(*γ, µ*) = argmin_*w*_ ℒ_train_(*w, γ, µ*), and for each feature space *i*, 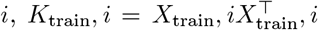 is the training kernel, and 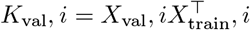 is the validation cross-kernel.

With this parameterization, hyperparameter random search is particularly efficient for multiple voxels. For each potential value of the kernel weights *γ* ∈ Δ_*m*−1_, the subproblem is a kernel ridge regression. Using the computational tricks described in Section 3.2, this kernel ridge regression is solved efficiently for multiple voxels and for multiple regularization parameters *µ*.

#### Dirichlet distribution

The kernel weights vector *γ* is defined on the simplex Δ_*m*−1_. To generate a candidate *γ* for random search, the Dirichlet distribution is used. The Dirichlet distribution is specifically defined on the simplex, and it is parametrized to prioritize different levels of sparsity. Its probability density function is parameterized by a set of concentration parameters 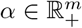 and reads 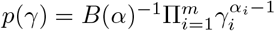, where *B*(*α*) is a normalization factor written with the gamma function *γ*. Because in the encoding model framework there is no prior reason to favor one feature space over another, the same value *α*_*i*_ is used for all dimensions. The value *α*_*i*_ = 1 for all *i* leads to a uniform distribution over the simplex. Smaller values increase the density near corners and edges of the simplex, leading to kernel weights that prioritize smaller subsets of kernels.

#### Full algorithm

The steps to implement hyperparameter random search are as follows. First, a set of candidates for *µ* is defined. These are most often specified on a unidimensional grid of logarithmically spaced values (*e*.*g*. {1, 10, 100}). Second, a set of candidates for *γ* is defined, sampled from the Dirichlet distribution. Third, for each candidate *γ*, the weighted-average kernel *K* = Σ_*i*_ *γ*_*i*_*K*_*i*_ is computed. With this single kernel, a kernel ridge regression is solved for all cross-validation splits, all regularizations *µ*, and all voxels. Fourth, the validation losses are averaged over splits, and their minimum is taken over regularizations *µ*. Finally, the hyperparameters *µ* and *γ* that minimize the average validation loss are selected on each voxel independently. (The entire algorithm is listed in pseudo-code in Appendix A.7.) Note that this algorithm is equivalent to the one described in [van de Wiel et al., 2021], except for the use of the Dirichlet distribution.

Hyperparameter random search is particularly efficient for large numbers of voxels, because the most costly computations can be factorized and reused on each voxel. This property is critical in voxelwise encoding models, where a separate model is fit independently on about 10^5^ voxels or more. However, hyperparameter random search has limited efficiency when the number of dimensions *m* (the number of feature spaces) increases. Indeed, the number of samples required to cover a *m*-dimensional space is of the order of 𝒪 (*e*^*m*^). Therefore, hyperparameter random search is not efficient when using large numbers of feature spaces. To address this issue, we propose a second method to optimize hyperparameters in multiple-kernel ridge regression.

### 3.6 Method 2: Hyperparameter gradient descent

Because hyperparameter random search has limited efficiency when the number of feature spaces is large, we propose a second method to solve banded ridge regression. The second method is hyperparameter gradient descent [Bengio, 2000]. Hyperparameter gradient descent is a method that iteratively improves hyperparameters. At each iteration, the update is based on the gradient of the cross-validation loss ℒ_val_ with respect to hyperparameters. Thus, the updated hyperparameters progressively converge toward the hyperparameters that minimize the cross-validation loss. The gradient of the cross-validation loss ℒ_val_ with respect to hyperparameters can be computed with implicit differentiation [Larsen et al., 1996; Chapelle et al., 2002; Foo et al., 2007], as described in details in Appendix A.4.

In the specific case of multiple-kernel ridge regression, the method is used to optimize *m* hyperparameters. In Section 3.5, the hyperparameters are parameterized with *γ* ∈ Δ_*m*−1_ and *µ* > 0, but this parameterization is inefficient for gradient descent. Indeed, gradient descent does not guarantee that the updated hyperparameters satisfy the constraints *γ* ∈ Δ_*m*−1_ and *µ* > 0. Thus, after each gradient descent update, an additional step is required to project the hyperparameters on their constrained subspace. Projecting *µ* on the positive subspace is computationally cheap, but projecting *γ* on the simplex Δ_*m*−1_ is more expensive. To avoid this cost, another parameterization is used here, defining *δ*_*i*_ = log(*γ*_*i*_*/µ*). Because *δ* ∈ R^*m*^ is unconstrained, no additional projection step is required. Furthermore, the logarithm improves the gradient conditioning, because *γ*_*i*_*/µ*typically spans multiple orders of magnitude. The training and validation loss functions are then defined as

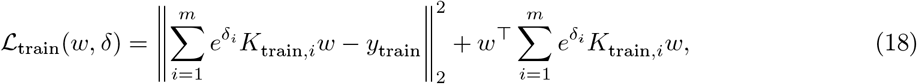

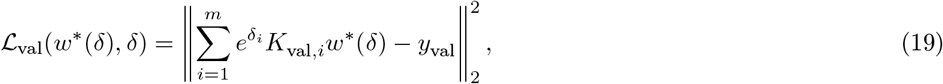

where *w*^*\**^(*δ*) = argmin_*w*_ ℒ_train_(*w, δ*), and for each feature space *i*, 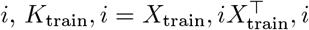 is the training kernel, and 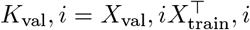 is the validation cross-kernel. Note that the optimization problem (19) is identical to (17), up to a reparametrization. See Appendix A.4 for the computation of the gradient of the cross-validation loss ℒ_val_ with respect to hyperparameters *δ*.

### 3.7 Computational complexity comparison

One straightforward way to evaluate different optimization methods is to compare them in terms of computational complexity. For example, one can estimate the computational cost of increasing the number of voxels or the number of feature spaces. Table 1 summarizes the computational complexity of the methods presented above for optimizing hyperparameters for multiple-kernel ridge regression: random search and gradient descent.

**Table 1:** Dominant computational complexities per iteration, for both hyperparameter optimization methods. The computational cost of the dominant computations changes when the size of the problem varies. The computational complexities give the order of change in computational cost when each dimension of the problem varies. Here, the different dimensions are the number of kernels *m* (*i*.*e*. the number of feature spaces), the number of training samples *n*_train_, the number of validation samples *n*_val_, the number of voxels *v*, the number of regularization candidates (for *µ*) *r*, and the number of cross-validation splits *s*. Importantly, the random-search diagonalization, which is the most expensive computation in the random-search method, is independent of the number of voxels *v*. On the contrary, the gradient-descent gradient computation is proportional to *v*. Therefore, gradient descent is fast for small numbers of voxels, but as the number of voxels increases, random search becomes more and more competitive compared to gradient descent.

| Random search |  |
| --- | --- |
| Kernel sum | $\mathcal{O}(mn_{\text{train}}^2)$ |
| Diagonalization | $\mathcal{O}(n_{\text{train}}^3 s)$ |
| Predictions | $\mathcal{O}(n_{\text{val}} n_{\text{train}} (n_{\text{train}} + v) r s)$ |
| Gradient descent |  |
| Lipschitz for $w$ (only once) | $\mathcal{O}(mn_{\text{train}}^3 s)$ |
| Gradient | $\mathcal{O}(mn_{\text{train}} (n_{\text{train}} + n_{\text{val}}) v s)$ |
| Lipschitz for $\delta$ | $\mathcal{O}(m^2 n_{\text{val}} v s)$ |

The most important difference between both proposed methods is the complexity with respect to the number of voxels *v*. In random search, the diagonalization is the most expensive operation, but it is done only once for all voxels. Therefore, the computational cost of random search remains almost the same for small or large numbers of voxels. On the contrary in gradient descent, the gradient computations are proportional to the number of voxels *v*. Therefore, the computational cost of gradient descent is low for small numbers of voxels, and high for large numbers of voxels. Comparing both methods, gradient descent is likely to be faster than random search for small numbers of voxels, and random search is likely to be faster than gradient descent for large numbers of voxels.

Another key parameter that affects computational complexity is the number of kernels *m* (*i*.*e*. the number of feature spaces in banded ridge regression). In the case of random search, the computational cost per iteration is barely affected by *m*, but the required number of iterations in a random search grows exponentially with *m*. In the case of gradient descent, the number of iterations required to reach convergence is invariant with respect to *m*, but the computation of each iteration is proportional to *m*. Comparing both methods, random search is likely to be faster than gradient descent for small numbers of feature spaces, and gradient descent is likely to be faster than random search for large numbers of feature spaces.

Note that in gradient descent, although the Lipschitz constant computation is inexpensive compared to the other steps, its complexity follows *m*^2^. Its cost might therefore become prohibitive for large numbers of feature spaces *m*, and an adaptive step-size strategy might become necessary (see for instance [Pedregosa, 2016]). In the example presented in Section 3.8, the Lipschitz constant was computed for up to *m* = 22 feature spaces, which is probably sufficient for most applications.

### 3.8 Banded ridge regression solver comparison

To demonstrate the computational efficiency of the proposed solvers, different banded ridge regression solvers were compared on three fMRI datasets. The number of time samples, feature spaces, cross-validation splits, and voxels of each dataset are detailed in Table 2. The stimuli used in each dataset were silent natural short clips watched with fixation (*“short-clips”* dataset) [Huth et al., 2012], natural stories listened with eyes closed (*“stories”* dataset) [Huth et al., 2016], and non-silent natural short films watched with eye tracking (*“short-films”* dataset) [Nunez-Elizalde et al., 2018]. Each dataset contained a number of manually engineered feature spaces, such as spatio-temporal wavelet transform magnitudes of visual stimuli [Nishimoto et al., 2011], hierarchical classification of objects present in the scenes [Huth et al., 2012], or word embeddings of speech content [Huth et al., 2016].

**Table 2:** Main dimensions of the three datasets used in our example. The dimensions include the number of training time samples *n*_train_ + *n*_val_, the number of testing time samples *n*_test_, the number of feature spaces *m*, the number of cross-validation splits *s*, and the number of voxels *v*. All datasets had a large number of voxels (*v* ≈ 10^5^). The largest number of feature spaces is *m* = 22 (see Appendix A.2 for a list of the 22 feature spaces).

| dataset | $n_{\text{train}} + n_{\text{val}}$ | $n_{\text{test}}$ | $m$ | $s$ | $v$ |
| --- | --- | --- | --- | --- | --- |
| short-clips | 3600 | 270 | 2 | 12 | 73221 |
| stories | 3737 | 291 | 4 | 10 | 73165 |
| short-films | 3572 | 807 | 22 | 12 | 85483 |

In all datasets, the number of features was larger than the number of time samples. Thus, all solvers used the multiple-kernel ridge regression formulation. A separate linear kernel was defined per feature space, and the different solvers described in Section 3.5, Section 3.6, and Appendix A.5 were compared.

#### Compared methods

In this example, nine different methods were empirically compared. The first three methods were used as baselines, and did not use banded ridge regression. The first baseline method used a ridge regression model fit on all feature spaces jointly, and with a hyperparameter *µ* shared across all voxels (as in [Huth et al., 2016]). The second baseline method used a ridge regression model fit on all feature spaces jointly, and with a different hyperparameter *µ* per voxel. In the third baseline, a separate ridge regression model was first fit on each feature space. Then, the best feature space was selected for each voxel independently, as measured by the average cross-validation loss.

The other six compared methods were fitting a banded ridge regression model, formulated as a multiple-kernel ridge regression. They only differed in the method used to solve the optimization problem. The hyperparameter gradient-descent was used with three variants, using either the direct gradient approximation (*“direct”*), the conjugate gradient approximation [Pedregosa, 2016] (*“conjugate”*), or the finite Neumann series approximation [Lorraine et al., 2019] (*“Neumann”*). The hyperparameter random search was used as described in Section 3.5. The proposed methods were also compared to the *Tikreg*^2^ implementation published previously by our group [Nunez-Elizalde et al., 2019], which explores hyperparameter space using Bayesian optimization of the average voxel performance (see more details in Appendix A.5). All methods were implemented on GPU, except the Tikreg implementation, which was only available on CPU. To give a fair comparison with the Tikreg implementation, the hyperparameter random search was also run on CPU.

#### Hyperparameter optimization

For the ridge regression models, the regularization hyperparameter *λ* was optimized over a grid-search with 20 values spaced logarithmically from 10^−5^ to 10^15^. This range was large enough to explore many different possible regularization strengths for each feature space. For hyperparameter random search, the Dirichlet concentration *α* alternated between three values 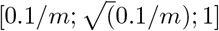 to explore both small and large subsets of kernels. A scaling by 1*/m* was used to get consistent kernel subset sizes over the different datasets. A grid of 20 values was used for *µ*, spaced logarithmically from 10^−5^ to 10^15^.

All gradient descent methods were initialized with the optimal regularization hyperparameter of the ridge regression baseline, computed on each voxel independently. For the conjugate gradient approximation, a decreasing precision *ε* was used with an exponential schedule [Pedregosa, 2016] going from 10^1^ to 10^−1^, for both subproblems (other precision schedules were tested and led to slower convergence). The other gradient-descent approximations used a single iteration of gradient descent to update the dual weights. The Neumann approximation was used with *k* = 5 (other values were tested and led to slower convergence).

As in previous examples, a leave-one-run-out cross-validation scheme was used, and the model was refit on the entire training dataset with the best hyperparameters. The solver comparison was performed on a Nvidia Titan X GPU (12 GB RAM). Because Tikreg was only available on CPU, the random-search solver was also run on CPU along with Tikreg, using a 8-core Intel CPU (3.4 GHz, 30 GB RAM).

#### Method evaluation

To evaluate the different methods, two criteria were used. The first criterion was the convergence of the validation loss ℒ_val_ averaged across cross-validation splits. The convergence of the validation loss over time is useful to compare the speed of the different methods to solve the optimization problem. The second criterion was the generalization *R*^2^ score on a test data set not used in model fitting. The generalization score is useful to verify that the changes in the validation loss are meaningful and not merely overfitting. The generalization score is also useful to compare banded ridge regression with the different baseline methods based on ridge regression. Note that in all three datasets, most voxels did not get any predictive power from the available feature spaces. Thus, the scores are averaged on a selection of best predicted voxels, selected using the top 10% generalization scores of ridge regression with shared regularization *µ*. Durations correspond to computations on all voxels.

## Results

The results of the method comparison are displayed in Figure 9. First, the Bayesian search implemented in Tikreg appeared several orders of magnitude slower than the proposed methods. This difference is not only due to the GPU implementation, because Tikreg’s Bayesian search was also slower than random search run on CPU. Bayesian search also sometimes generalized poorly (short-clips and short-films datasets). This poor generalization can be explained by the optimization being stopped early due to computational constraints. Another possible explanation is that Bayesian search is solving an approximation of the problem, while the proposed methods solve the correct problem (see Appendix A.5).

**Figure 9:**
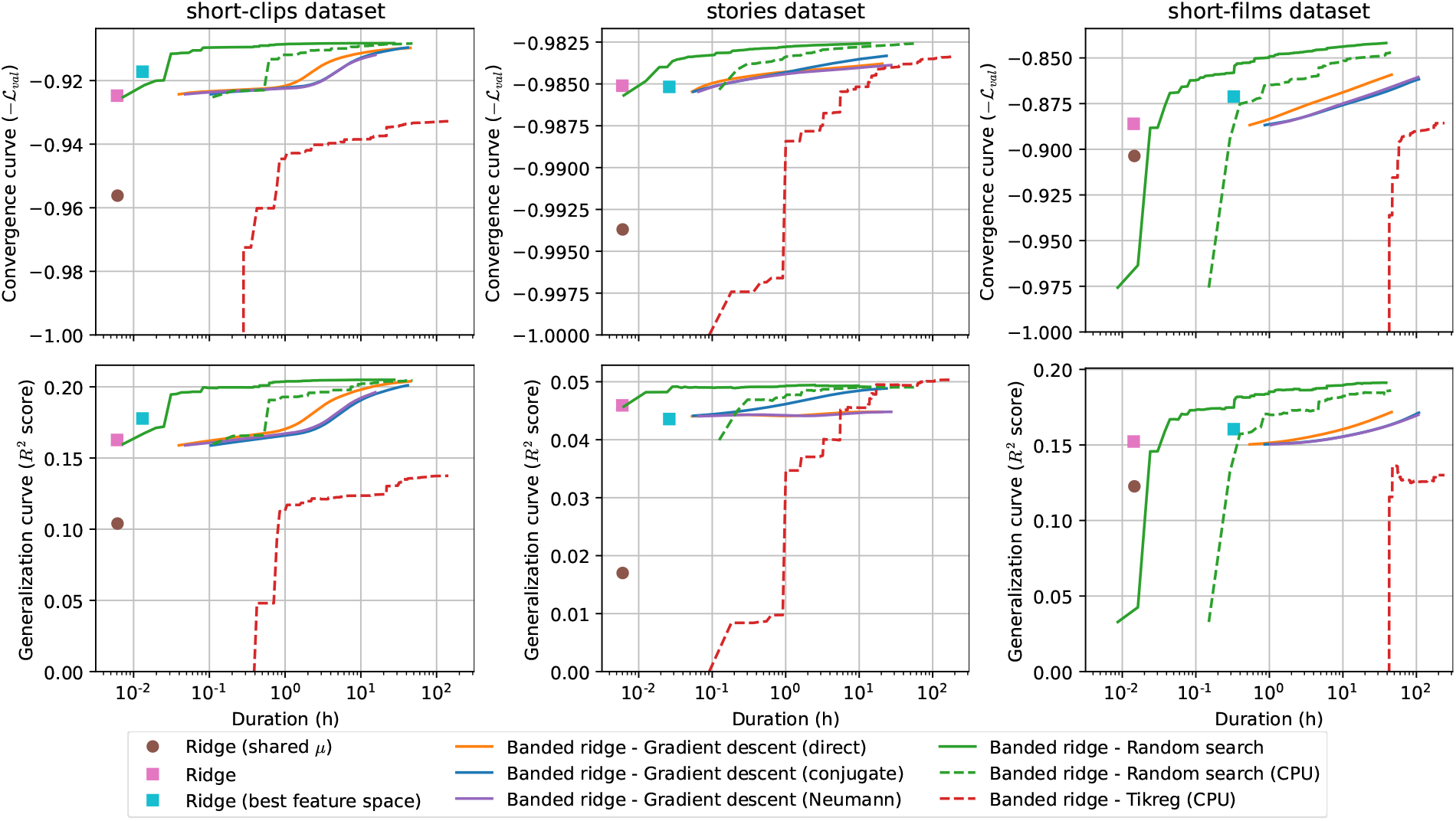
Convergence and generalization of different banded ridge regression solvers, on three different fMRI datasets. All x-axes indicate duration in logarithmic scale. All methods were run on GPU, except when plotted with dashed lines. **(Top row)** Convergence over time, as measured with the negative validation loss (−ℒ_val_) (higher is better), averaged over splits and voxels. Convergence over time measures the speed of convergence of the different methods used to fit banded ridge regression. Random search is the fastest method on all three datasets, thanks to its efficient factorization over voxels. In particular, it is more efficient than Tikreg’s Bayesian search by several orders of magnitude, even when performed on CPU. **(Bottom row)** Generalization performances over time, as measured by the *R*^2^ score computed on a test set (higher is better), averaged over voxels. The generalization score is useful to verify that the changes in the validation loss are meaningful and not merely overfitting. The generalization score is also useful to compare banded ridge regression with different baseline models. Overall, banded ridge regression reaches higher generalization scores than the ridge regression baselines. The proposed solvers also reach higher generalization scores than Tikreg’s Bayesian search, given a reasonable time budget.

Among the proposed methods, random search was faster than gradient descent, even for a medium number of feature spaces *m* = 22 (short-films dataset). This might be attributed to the factorization over voxels of random search, which is critical for large numbers of voxels *v* ≈ 10^5^. This is especially important as the number of voxels can grow even larger for higher resolution fMRI (up to 10^7^ in recent datasets [Feinberg et al., 2018]. However, it is expected that gradient descent outperforms random search for larger numbers of feature spaces.

The comparison of the three gradient-descent approximations was not conclusive. On the stories dataset, both direct and Neumann approximations did not generalize as well as other solvers. On the short-clips and short-films datasets, the direct gradient was faster, though one would need further convergence to compare generalization. Importantly, the direct gradient approximation performed well in two out of three datasets. This result is surprising, as the direct gradient is a strong approximation with no guarantee to converge toward a correct solution.

Finally, banded ridge regression generalized better than all ridge regression baselines. This improvement can be explained by the feature-space selection mechanism of banded ridge regression, which effectively removes non-predictive and redundant feature spaces, while still allowing multiple feature spaces to be complementary.

### 3.9 Recommended strategy

Based on the empirical solver comparison, we recommend using random search to efficiently solve banded ridge regression on large numbers of voxels. However, the convergence of random search is fast in the first iterations but slower afterward. Moreover, the Dirichlet sampling used in random search introduces a prior which biases the hyperparameter optimization. To find more precise feature-space selection and fix the prior bias, one can use gradient descent to refine the solution on a selection of best performing voxels. To do so, the hyperparameters selected by random search can be used to initialize gradient descent. This refinement can lead to a better feature-space selection, and thus improve the variance decomposition interpretation. To limit the computational cost of the gradient-descent refinement, the refinement can be performed only on a selection of best-predicted voxels, as measured by the cross-validation loss.

### 3.10 Python package

All methods described in this paper are implemented in an open-source Python package called *Himalaya*. All methods are both available through a functional API and through a class API compatible with scikit-learn [Pedregosa et al., 2011]. Moreover, to use either CPU or GPU resources, the package can be used seamlessly with different computational backends based on *Numpy* [Harris et al., 2020], *Pytorch* [Paszke et al., 2019], or *Cupy* [Nishino and Loomis, 2017]. Finally, the package includes extensive documentation, rigorous unit testing (including scikit-learn’s estimator checks), and a gallery of examples to facilitate its use.

## Conclusion

Banded ridge regression is a natural extension to ridge regression that takes into account any predefined group structure in the features. In neuroimaging studies using encoding models, a group structure is naturally defined when different feature spaces correspond to different hypotheses or different representations of the stimuli and tasks. Banded ridge regression is able to adapt regularization to this structure and to learn optimal scalings for each feature space. These optimal scalings are learned by cross-validation, a principled method that focuses on maximizing generalization to unseen datasets. Banded ridge regression is thus a robust model that adapts regularization to predefined group structures in the features to maximize generalization in linear encoding models.

In this paper, we argue that the feature-space selection mechanism of banded ridge regression partly explains its good performance. By learning optimal feature-space scalings during cross-validation, banded ridge regression is able to ignore some non-predictive or redundant feature spaces. Ignoring these feature spaces reduces the tendency to overfit, and focuses prediction on feature spaces that are most likely to generalize. Moreover, this feature-space selection improves interpretability, because it reduces the number of feature spaces that contribute to model predictions. This feature-space selection is especially interesting when using naturalistic stimuli that are often best modeled using highly correlated feature spaces. Because banded ridge regression disentangles correlated feature spaces it clarifies which specific feature spaces best predict brain activity.

In this paper we also relate feature-space selection to a large literature of sparsity-inducing linear regression methods. In algorithms ranging from automatic relevance determination to multiple-kernel learning and the group lasso, the selection of feature spaces (or features) has been shown multiple times to lead to better generalization and better interpretation. In the encoding model framework, generalization to unseen data is a powerful way to validate the model and to improve reproducibility. Generalization performance also serves as a guide for model selection and interpretation. By maximizing both interpretation and generalization, banded ridge regression is a powerful tool for data-driven analysis of rich naturalistic experiments in neuroimaging encoding models.

Finally, we demonstrate how the computational cost of banded ridge regression can be largely reduced compared to the original implementation. Our proposed methods use principled algorithms such as hyperparameter random search and hyperparameter gradient descent. In particular, we show how to use these algorithms to fit a separate banded ridge regression on each voxel independently, scaling efficiently to large numbers of voxels. With the advent of layer-specific 7T BOLD fMRI at sub-millimeter resolution, the number of voxels is expected to increase by one or two orders of magnitude compared to the 3T fMRI datasets used in this work. Using efficient algorithms implemented on GPU is thus critical to be able to scale the analysis to the new orders of magnitude of data to come in the future. To facilitate dissemination, all algorithm implementations are released in an open-source, GPU-compatible Python package called *Himalaya*^3^.

## Acknowledgments

This work was supported by grants from the National Eye Institute (R01-EY031455), the National Science Foundation (Nat-1912373), the Office of Naval Research (N00014-20-1-2002), the National Institutes of Health (B-U01EB02), the Weill Neurohub at University of California, Berkeley, and internal funds from University of California, Berkeley.

This work was also made possible thanks to the scientific Python ecosystem, including *Numpy* [Harris et al., 2020], *Scipy* [Virtanen et al., 2020], *Matplotlib* [Hunter, 2007], *Scikit-learn* [Pedregosa et al., 2011], *Pycortex* [Gao et al., 2015], *Pytorch* [Paszke et al., 2019], and *Cupy* [Nishino and Loomis, 2017].

We also thank Bin Yu, Fabian Pedregosa, Matteo Visconti di Oleggio Castello, Storm Slivkoff, and members of the Gallant lab for fruitful discussions and comments.

## Contributions

Conceptualization, TDT, ME, and AONE; Software, TDT and ME; Methodology, TDT; Formal analysis, TDT and ME; Investigation, TDT; Visualization, TDT; Writing - Original Draft, TDT; Writing - Review & Editing, TDT, ME, AONE, and JLG; Funding acquisition, JLG.

## Code availability statement

All methods described in this paper are implemented in an open-source Python package called *Himalaya* (https://github.com/gallantlab/himalaya).

## Ethics statement

Data was acquired on human subjects during previous studies ([Huth et al., 2012, 2016; Nunez-Elizalde et al., 2018]. The experimental protocols were approved by the Committee for the Protection of Human Subjects at University of California, Berkeley. All subjects were healthy, had normal hearing, and had normal or corrected-to-normal vision. Written informed consent was obtained from all subjects. The authors declare no competing financial interests.

## A Supplementary materials

### A.1 Quantifying feature-space selection with the effective rank

In Section 2.4 and Section 2.7, two examples are proposed to demonstrate the feature-space selection mechanism in banded ridge regression. For the sake of demonstration, these examples use the *effective rank* to quantify the number of feature spaces effectively used in the model. This section explains why the effective rank metric was chosen, and compares alternative definitions of the effective rank.

To quantify the number of feature spaces effectively used in the model, one candidate metric is to count the number of feature spaces with a non-zero effect on the prediction. However, because the hyperparameters optimized by banded ridge regression never reach infinity, the coefficients *b*^*\**^ are never exactly zero, even for feature spaces that are effectively ignored. For this reason, all feature spaces have non-zero values in the variance decomposition, and one cannot simply count the number of non-zero values. To fix this issue, it is possible to define a positive threshold, and to count the number of values above the threshold. Yet, this threshold would have an arbitrary value, and would artificially separate values that are slightly above from values that are slightly below, leading to a discontinuity in the metric. Instead, a better metric of the number of feature spaces effectively used in the model would vary continuously as some feature space contribution goes to zero.

To address this issue, we leverage a metric to compute the rank of a linear system. The rank of a linear system is the number of independent equations in the system, and corresponds to the number of non-zero eigenvalues *λ*_*i*_. The rank is thus a discontinuous function of the eigenvalues. To remove the discontinuity, the rank can be approximated by the *effective rank*. Several effective rank definitions exist, such as *e* [Roy and Vetterli, 2007], *r*_0_ [Bartlett et al., 2020], *R*_0_ [Langeberg et al., 2019; Bartlett et al., 2020], and *h* (proposed in this work). Assuming the eigenvalues are non-negative and sorted in decreasing order (*λ*_0_ ≥ *λ*_1_ ≥ … ≥ *λ*_*m*−1_ ≥ 0), these four metrics are defined as

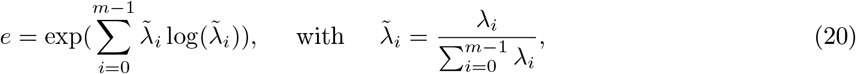

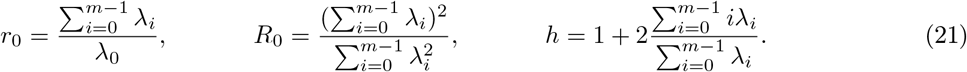

All four metrics (*e, r*_0_, *R*_0_, and *h*) have similar useful properties. They are continuous functions of the *m* eigenvalues, with values in the interval [1, *m*]. Because they are continuous, they do not require defining any threshold. They are also not affected by additional unused dimensions (dimensions *i* with *λ*_*i*_ = 0). They are equal to *k* when *k* eigenvalues are equal (and non-zero) and all the other eigenvalues are equal to zero. The four metrics only differ in the way these integer values are interpolated. To describe how they interpolate these integer values, Figure 10 presents the metric values on different 2D and 3D spaces.

**Figure 10:**
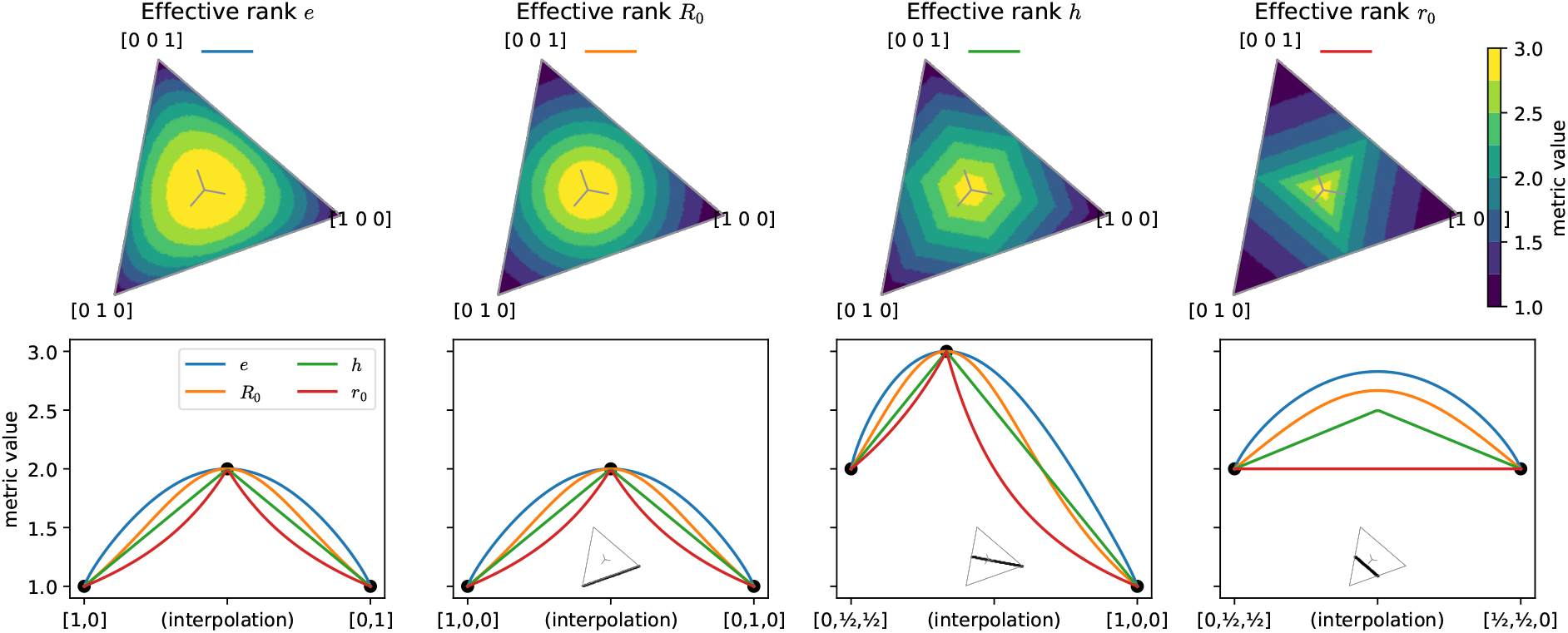
Comparison of four effective rank metrics. Effective rank metrics are useful to quantify the number of feature spaces used in banded ridge regression. **(Top)** Values of the four metrics on the simplex. The variance decompositions are defined on the simplex Δ_*m*−1_, the space of vectors in dimension *m* with positive values that sum to one. With *m* = 3, the simplex is a 2-dimensional surface in the shape of a triangle. All four effective rank metrics are equal to 1 at the triangle corners, to 2 at the edge midpoints, and to 3 at the center. The four metrics differ in the way they interpolate these integer values. The color mapping is discrete to highlight the structures of the different metrics. **(Bottom)** Values of the four metrics on different 1-dimensional trajectories, linearly interpolating between two points in dimension 2 or 3. All four metrics are continuous, and equal to *k* when the values are equally split over *k* dimensions (depicted by the black dots). Moreover, adding unused dimensions (second versus first panel) does not alter the metric values. All four metrics are thus reasonable choices for quantifying the number of feature spaces used in banded ridge regression. The definition *e* was used throughout this paper for its intuitive link to entropy. features; Nishimoto et al. [2011]).

To use an effective rank metric to quantify the number of feature spaces effectively used in the model, the following procedure is used. First, the variance decomposition *ρ* ∈ ℝ ^*m*^ is computed, for example using the product measure as described in Section 1.6. Then, negative values are clipped to zero, and an effective rank metric is computed on the vector *ρ*. Although all four metrics give similar results, the definition *e* was used throughout this paper for its intuitive link to the entropy *H* = Σ _*i*_ *λ*_*i*_ log(*λ*_*i*_) [Roy and Vetterli, 2007; Araki and Lieb, 2002; Bach, 2022]. Because the effective rank estimates the number of feature spaces effectively used over *m* features spaces, it is noted 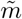.

### A.2 List of the 22 feature spaces from the short-films dataset

In Section 2.6 and Section 3.8, banded ridge regression is fit on 22 features spaces from the “short-films” dataset [Nunez-Elizalde et al., 2018]. Here, a short description of the 22 feature spaces is given.

- Stimulus motion energy: spatiotemporal Gabor filters computed on the stimulus luminosity (11,845 features; Nishimoto et al. [2011]).
- Retinotopic motion energy: spatiotemporal Gabor filters, computed on the stimulus luminosity, corrected for eye movements (11,845 features).
- Eye position: cubic polynomial expansion of the eye position during movie watching (36 features).
- Visual semantic: semantic labels of objects and actions present in the scene [Huth et al., 2012], projected in a semantic space (985 features; Huth et al. [2016]).
- Visual thematics (5 feature spaces): categorization of verb classes into categories based on the type of events described. VerbNet (85 features; Kipper et al. [2006]) was used to construct a thematic roles feature space (36 features), a selective restrictions feature space (38 features) and their interaction (71 features), and a verb categories feature space (101 features).
- Spectrogram: spectral power density of the sound wave (2388 features).
- Wavenet: Auditory features related to musical instruments extracted using a pretrained convolutional neural network (512 features; Oord et al. [2016]).
- Word rate: cubic polynomial expansion of word rate (13 features).
- Speech syntax position: part-of-speech tags of each word in the sentences, extracted using a pre-trained neural network (12 features; Andor et al. [2016]).
- Speech syntax labels: word dependencies in the sentence, extracted using a pre-trained neural network (43 features; Andor et al. [2016]).
- Speech semantic: spoken words projected in a semantic space (985 features; Huth et al. [2016]).
- Speech thematics (7 feature spaces): categorization of verb classes into categories based on the type of events described. VerbNet [Kipper et al., 2006] labels were used to construct a thematic role feature space (39 features), a verb class feature space (274 features), and a verb category feature space (102 features). Four thematic roles (verb, actor, recipient, place) were further decomposed by projecting words in a semantic space (985 feature each).

### A.3 Banded ridge regression is equivalent to multiple-kernel ridge regression

As stated in Section 2.8, banded ridge regression is equivalent to multiple-kernel ridge regression, up to a subtle difference. This subsection first demonstrates the equivalence, then discusses the source and implications of the difference. Note that a deep understanding of the difference is not critical to the rest of the paper.

#### Equivalence

To reformulate banded ridge regression as a multiple-kernel ridge regression, one can note that banded ridge regression is equivalent to a ridge regression with weighted feature spaces 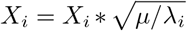 (see Section 2.2). This ridge regression is in turn equivalent to a kernel ridge regression [Saunders et al., 1998] (see Section 3.2) with a linear kernel 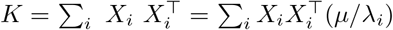. Writing 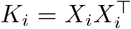 and *γ*_*i*_ = *µ/λ*_*i*_, the model becomes a multiple-kernel ridge regression

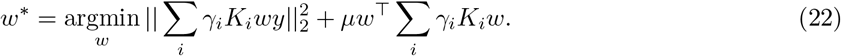

Multiple-kernel ridge regression corresponds to a kernel ridge regression with a weighted kernel *K* = Σ _*i*_ *γ*_*i*_*K*_*i*_ and a regularization hyperparameter *µ*. Because *µ* is arbitrary in the change of variables, one can choose it such that Σ _*i*_ *γ*_*i*_ =Σ _*i*_ *µ/λ=*1, which leads to *µ =* (Σ_*i*_1/ λ _*i*_)^−1^. In this way, is restricted to a subspace called the simplex 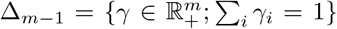 which corresponds to the standard formulation in the multiple-kernel learning literature [Gönen and Alpaydin, 2011]. Banded ridge regression is thus equivalent to multiple-kernel ridge regression. In Section 3.5, this equivalence is used to solve banded ridge regression efficiently when the number of features is larger than the number of samples.

#### Key difference

The equivalence between banded ridge regression and multiple-kernel ridge regression hides a subtle difference in the way hyperparameters are optimized. In banded ridge regression, all hyperparameters 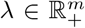 are optimized through cross-validation. Thus, in the equivalent multiple-kernel ridge regression, both the kernel weights *γ* ∈ Δ_*m*−1_ and the regularization *µ* > 0 must be optimized through cross-validation. On the contrary, in the multiple-kernel learning literature, the kernel weights *γ* are usually learned within the training set [Gönen and Alpaydin, 2011], and only *µ* is learned through cross-validation. Some rare papers have proposed learning kernel weights through cross-validation [Tsuda et al., 2004; Keerthi et al., 2007; Klatzer and Pock, 2015], but this practice is not common in the multiple-kernel learning literature. This difference is important, and is further discussed in the rest of the subsection. To highlight the difference between the two strategies, the following notations is used:

- **MKRR-cv** is a multiple-kernel ridge regression with kernel weights learned over **c**ross-**v**alidation.
- **MKRR-in** is a multiple-kernel ridge regression with kernel weights learned with**in** the training set.

Using these notations, banded ridge regression is equivalent to MKRR-cv, whereas the standard formulation in multiple kernel learning is MKRR-in. Additionally, the squared group lasso is equivalent to MKRR-in [Bach, 2008; Rakotomamonjy et al., 2008].

#### Group scalings

To understand the difference between MKRR-cv and MKRR-in, it is useful to consider the concept of group scalings. When feature spaces are unbalanced (for example in terms of number of features), the regularization can have a different strength on each feature space. To fix this issue, multiple studies have proposed that the regularization strength on each feature space could be evened out by adding group scalings to the regularization. In the group lasso literature, group scalings are positive weights *d*_*i*_ defined for each feature space *i*, which creates a scaled *L*_2,1_ norm Σ_*i*_ *d*_*i*_||*b*_*i*_||_2_. Similarly, in the multiple kernel learning literature, group scalings modify the optimization by constraining kernel weights to a scaled simplex 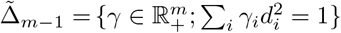 [Bach, 2008]. Note that group scalings are defined additionally to the kernel weights Because feature spaces can be unbalanced in different ways, many group scalings have been proposed in the group lasso and multiple-kernel learning literature: the square root of the number of features [Yuan and Lin, 2006], the kernel trace [Lanckriet et al., 2004], the square root of the kernel trace [Bach et al., 2004], the kernel empirical rank [Bach et al., 2005], or more sophisticated group scalings (*e*.*g*. [Bach, 2008; Obozinski et al., 2011; Pan and Zhao, 2016]). However, no group scaling emerged as being superior to the others. In MKRR-in, it is thus necessary to test different group scalings, and select the best one through cross-validation. On the contrary, in MKRR-cv the kernel weights *γ* ∈ Δ_*m*−1_ are learned by cross-validation. Thus, group scalings would have no effect on MKRR-cv. In other words, MKRR-cv is equivalent to using MKRR-in with group scalings optimized via cross-validation.

#### Summary

To summarize this subsection, MKRR-cv and MKRR-in are two variants of multiple-kernel ridge regression. In MKRR-cv, the kernel weights are learned using cross-validation, whereas in MKRR-in, the kernel weights are learned within the training set. Banded ridge regression is equivalent to MKRR-cv, whereas the squared group lasso is equivalent to MKRR-in. In MKRR-in, one can add group scalings to help balance the feature spaces, but the choice of group scaling needs to be cross-validated. In MKRR-cv, group scalings are less important because optimal kernel weights are already learned through cross-validation.

### A.4 Derivation of the hyperparameter gradient

In Section 3.6, hyperparameter gradient descent is used to optimize hyperparameters in multiple-kernel ridge regression. Hyperparameter gradient descent is a method that iteratively improves hyperparameters. At each iteration, the update is based on the gradient of the cross-validation loss *L*_val_ with respect to hyperparameters *δ*. Here, the computation of this gradient is derived using the implicit differention framework.

As explained in Section 3.6, multiple-kernel ridge regression is reparameterized with hyperparameters *δ* ∈ ℝ^*m*^. The training and validation loss functions are defined as

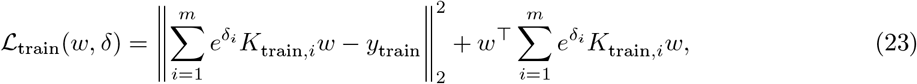

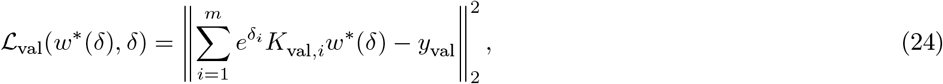

where *w*^*\**^(*δ*) = argmin_*w*_ ℒ_train_(*w, δ*), and for each feature space 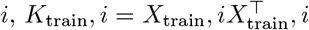 is the training kernel, and 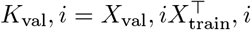*i* is the validation cross-kernel.

#### Implicit differentiation framework

At each iteration, gradient descent updates the hyperparameters *δ* in the direction of the gradient of the cross-validation loss. The cross-validation loss ℒ_val_(*w*^*\**^(*δ*), *δ*) has two dependencies in *δ*, one *direct*, and one *indirect* through the dual weights *w*^*\**^(*δ*). The gradient of ℒ_val_ with respect to *δ* is thus the sum of the direct gradient 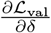 and the indirect gradient 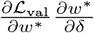. In the indirect gradient, the Jacobian 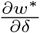 can be computed with *implicit differentiation* [Larsen et al., 1996; Chapelle et al., 2002; Foo et al., 2007], which leads to the full gradient expression

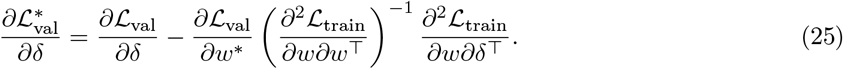

In this expression, the most expensive term to compute is the inverse Hessian 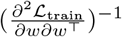 To reduce this computational bottleneck, three separate approximations are considered. The first approximation, proposed in [Pedregosa, 2016], uses a conjugate gradient method to solve 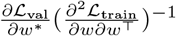 up to a limited precision. The second approximation, proposed in [Lorraine et al., 2019], uses Neumann series to estimate the inverse Hessian with 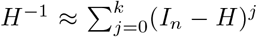 with typically *k* = 5. The third approximation is more aggressive, and approximates the full gradient with the direct gradient 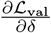 This last approximation is equivalent to assuming the inverse Hessian to be 0. In many applications, this approximation is not available because the direct gradient is equal to zero [Lorraine et al., 2019]. However, the direct gradient is not equal to zero in multiple-kernel ridge regression, so it can be used to approximate the full gradient.

#### Application to multiple-kernel ridge regression

The implicit differentiation framework can be immediately applied to multiple-kernel ridge regression. Yet, some improvements can be done to maximize efficiency. For example, the gradient of the training loss at optimum 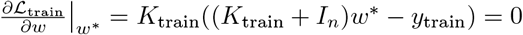 can be simplified to (*K*_train_ + *I*_*n*_)*w*^*\**^ − *y*_train_ = 0 by using the gradient with respect to the scalar product ⟨*x, y*⟩_*K*_ = *x*^⊤^*Ky*. This change improves gradient conditioning, and leads to the same solutions when the kernel has full rank. In addition, this change largely simplifies the following derivative expressions and decreases their computational cost

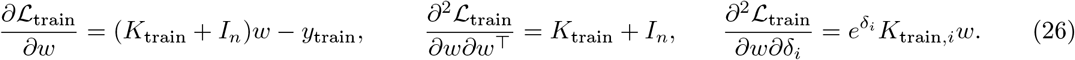

The other derivative expressions are not affected by this change, and are equal to

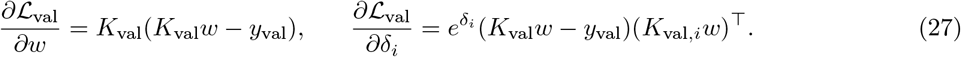

Once the gradient 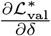 is computed, gradient descent improves the hyperparameters *δ* with the update 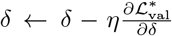 where *η* is called the step size. To estimate the optimal step size, one could use the Lipschitz constant of 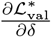 but that is hard to estimate [Pedregosa, 2016]. Fortunately, the Lipschitz constant of the direct gradient 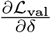 can be estimated. Noting 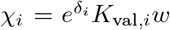, the validation loss reads 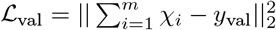 Using 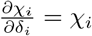 the gradient and Hessian are derived as

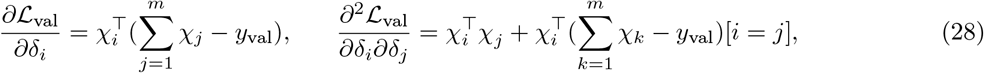

where [*i* = *j*] is equal to 1 if *i* = *j* and 0 otherwise. The local Lipschitz constant *L* is given by the maximum eigenvalue of the Hessian, which can be estimated with power iteration. In our experiments, a step-size *η* = 1*/L* was used, not only for the direct gradient, but also for the full gradient approximations. Several adaptive step-size strategies were also explored, but none were computationally as efficient as using the estimated Lipschitz constant of the direct gradient. The entire algorithm is listed in pseudo-code in Appendix A.7.

### A.5 Hyperparameter Bayesian search

In Section 3.5 and Section 3.6, two methods are proposed to solved banded ridge regression. Another method proposed in the past to solve banded ridge regression is hyperparameter Bayesian search [Nunez-Elizalde et al., 2019]. Hyperparameter Bayesian search [Mockus et al., 1978] is a method similar to hyperparameter random search. Both search methods compare a number of hyperparameter candidates and select the candidates with the lowest cross-validation error. The specificity of Bayesian search is that each candidate is selected conditionally to the performances of previously tested candidates. The *Tikreg* Python package [Nunez-Elizalde et al., 2019] published previously by our group implements Bayesian search to solve banded ridge regression. Specifically, Tikreg implements a Bayesian search using trees of Parzen estimators [Bergstra et al., 2011] through the Hyperopt Python package [Bergstra et al., 2013].

A major limitation of hyperparameter Bayesian search is that it cannot optimize multiple losses simultaneously. For this reason, Tikreg cannot optimize a separate loss per voxel. Instead, Tikreg optimizes the *average* validation loss over all voxels. This approximation introduces a bias towards the preferred hyperparameters of the voxel majority. To mitigate this bias, Tikreg later selects the best hyperparameters per voxel out of all candidates explored by Bayesian search. However, this mitigation will only be successful if Bayesian search explores the hyperparameter space broadly and does not find its optimum quickly.

Another limitation of Tikreg’s implementation of Bayesian search is that it does not use all computational tricks to solve the ridge regression subproblems. As described in Section 3.1, ridge regression can be solved efficiently by reusing computations for multiple regularization hyperparameters. To leverage this computational trick in random search (Section 3.5), hyperparameters are separated into *γ* ∈ Δ_*m*−1_ and *µ* > 0. For each randomly-sampled candidate *γ*, a ridge regression is solved efficiently for a large number of *µ*. This factorization is key to explore hyperparameter space efficiently, but is not present in Tikreg’s implementation.

### A.6 Algorithms for weighted-kernel ridge regression

To solve multiple-kernel ridge regression, the hyperparameter gradient descent solver updates the kernel weights *δ* ∈ R^*m*^ at each iteration. Then, at each iteration, the regression weights *w* must be optimized for the current (fixed) kernel weights *δ* ∈ ℝ^*m*^. We call this sub-problem *weighted-kernel ridge regression*. Specifically, for a *fixed δ* ∈ ℝ^*m*^, weighted-kernel ridge regression is defined as:

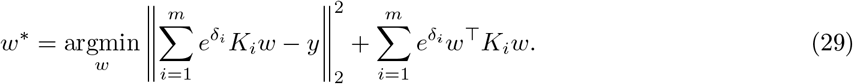

This section describes how to solve this sub-problem.

This problem has a closed formed solution *w*^*\**^ = (*K* + *I*_*n*_)^−1^*y*, where *K* is the kernel weighted sum 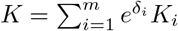. However, if different voxels have different kernel weights *δ*, this closed-form solution is not efficient, as it requries computing *K* and inverting (*K* + *I*_*n*_)^−1^ for each different kernel weights *δ*. During hyperparameter gradient descent, a precise solution of the weighted-kernel ridge regression sub-problem is not necessary, and an approximation is sufficient [Pedregosa, 2016]. Thus, an approximate solution can be obtained with gradient descent (Algorithm 1), conjugate gradient descent (Algorithm 3), or even finite Neumann series (Algorithm 2). (Neumann series approximates 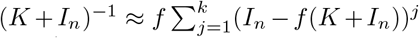 where the factor 0 < *f* < 1 needs to be tuned to make the series converge.) All three algorithms are presented here for a single voxel, yet they can be straightforwardly extended to multiple voxels using matrices and tensors.

Interestingly, another sub-problem of hyperparameter gradient descent can be solved as a weighted-kernel ridge regression. During hyperparameter gradient descent, the gradient computation requires computing 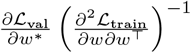 in multiple-kernel ridge regression, this term is equal to *K*_val_(*K*_val_*w* − *y*_val_) (*K*_train_+*I*_*n*_)^-1^.This expression can be considered as the solution of equation (29), with *y* = *K*_val_(*K*_val_*w* − *y*_val_). Therefore, the three approximate solvers of weighted-kernel ridge regression presented in this section can also be used to compute this element of the hyperparameter gradient computation.

In the example presented in Section 3.8, the ridge dual weights were updated with Algorithm 1 because Algorithm 2 was too coarse and Algorithm 3 led to oscillations in the convergence curves. For the hyperparameter gradient term, because the term did not require a precise approximation, all three algorithms (Algorithm 1, Algorithm 3, and Algorithm 2) were tested and compared.

#### Algorithm 1

Weighted-kernel ridge regression solver - Gradient descent

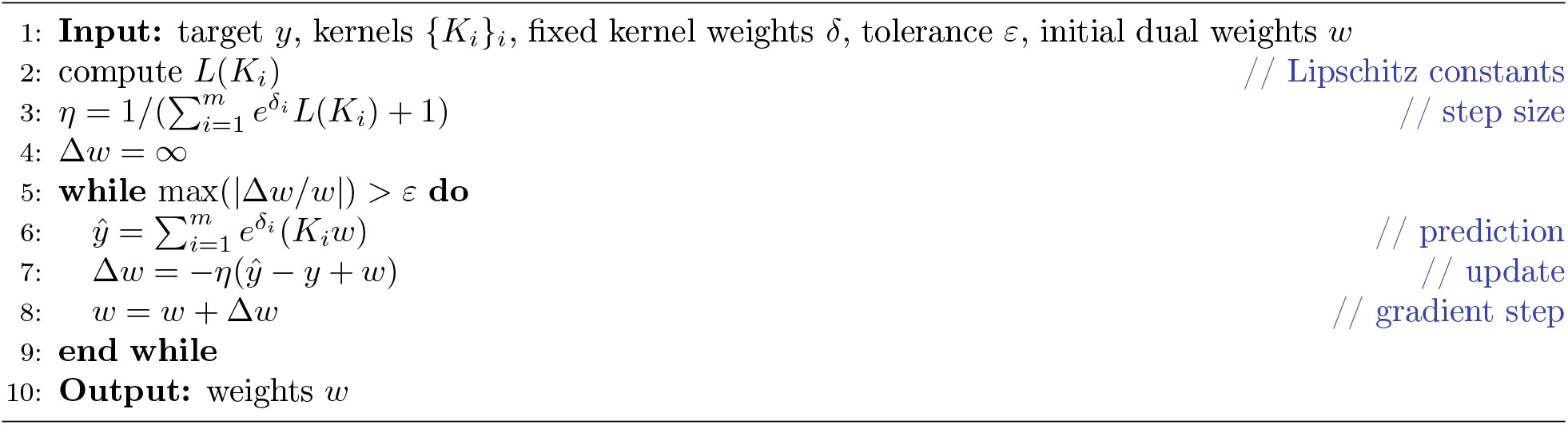

#### Algorithm 2

Weighted-kernel ridge regression solver - Neumann series

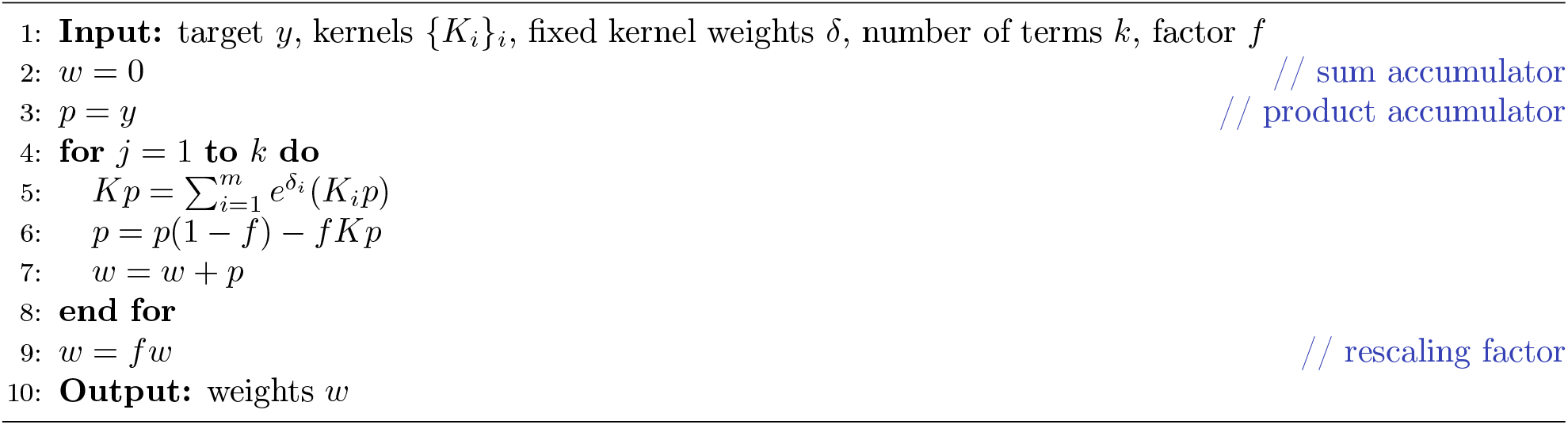

#### Algorithm 3

Weighted-kernel ridge regression solver - Conjugate gradient

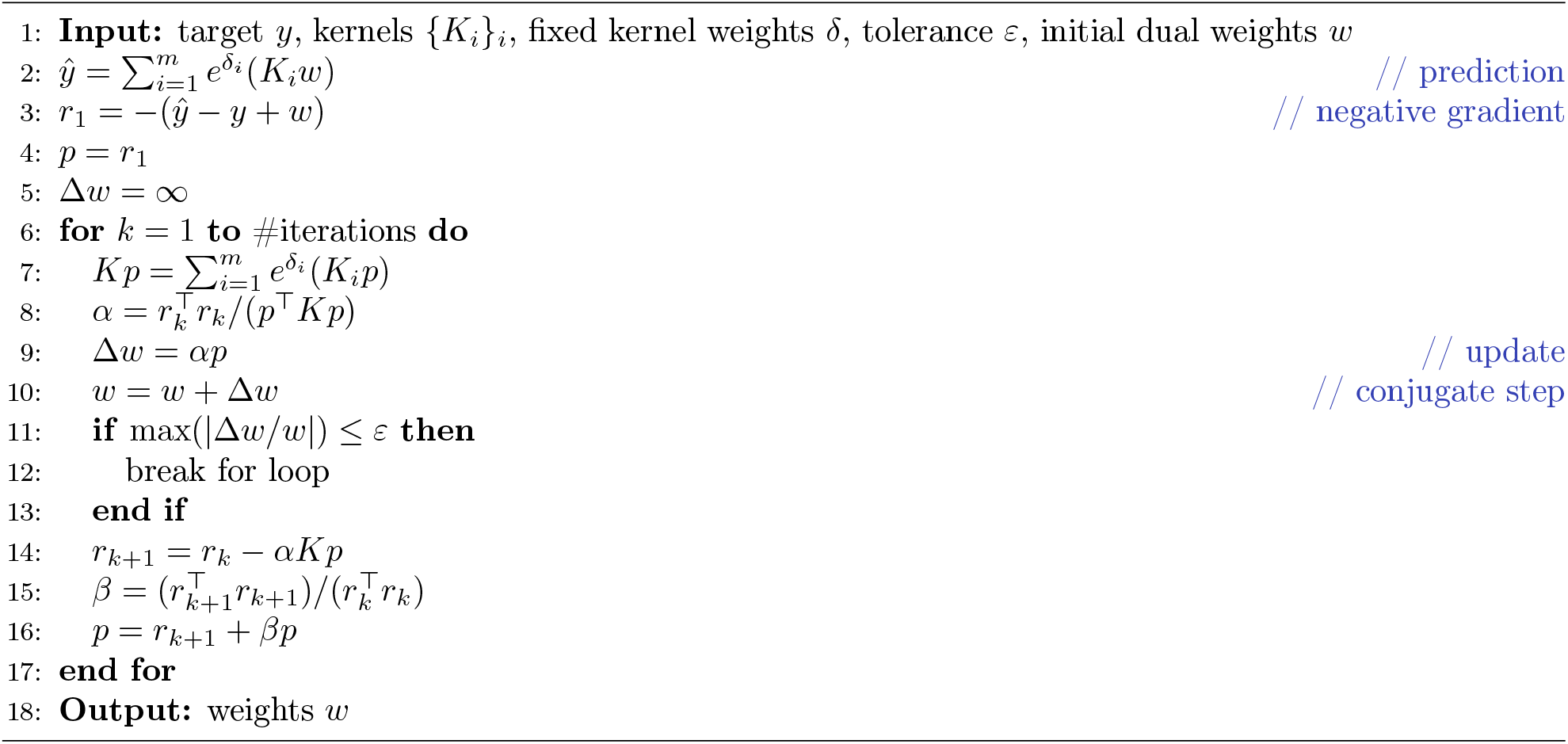

### A.7 Algorithms for multiple-kernel ridge regression

This section describes algorithms to solve multiple-kernel ridge regression, optimizing *δ* ∈ ℝ^*m*^ over the validation loss:

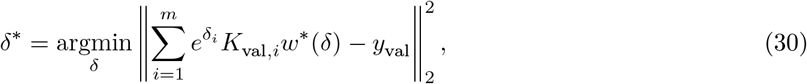

where *w*^*\**^(*δ*) is the solution of the weighted-kernel ridge regression (29). This problem can be either solved with hyperparameter random search (Algorithm 4) or with hyperparameter gradient descent (Algorithm 5).

#### Algorithm 4

Multiple-kernel ridge regression solver - Hyperparameter random search

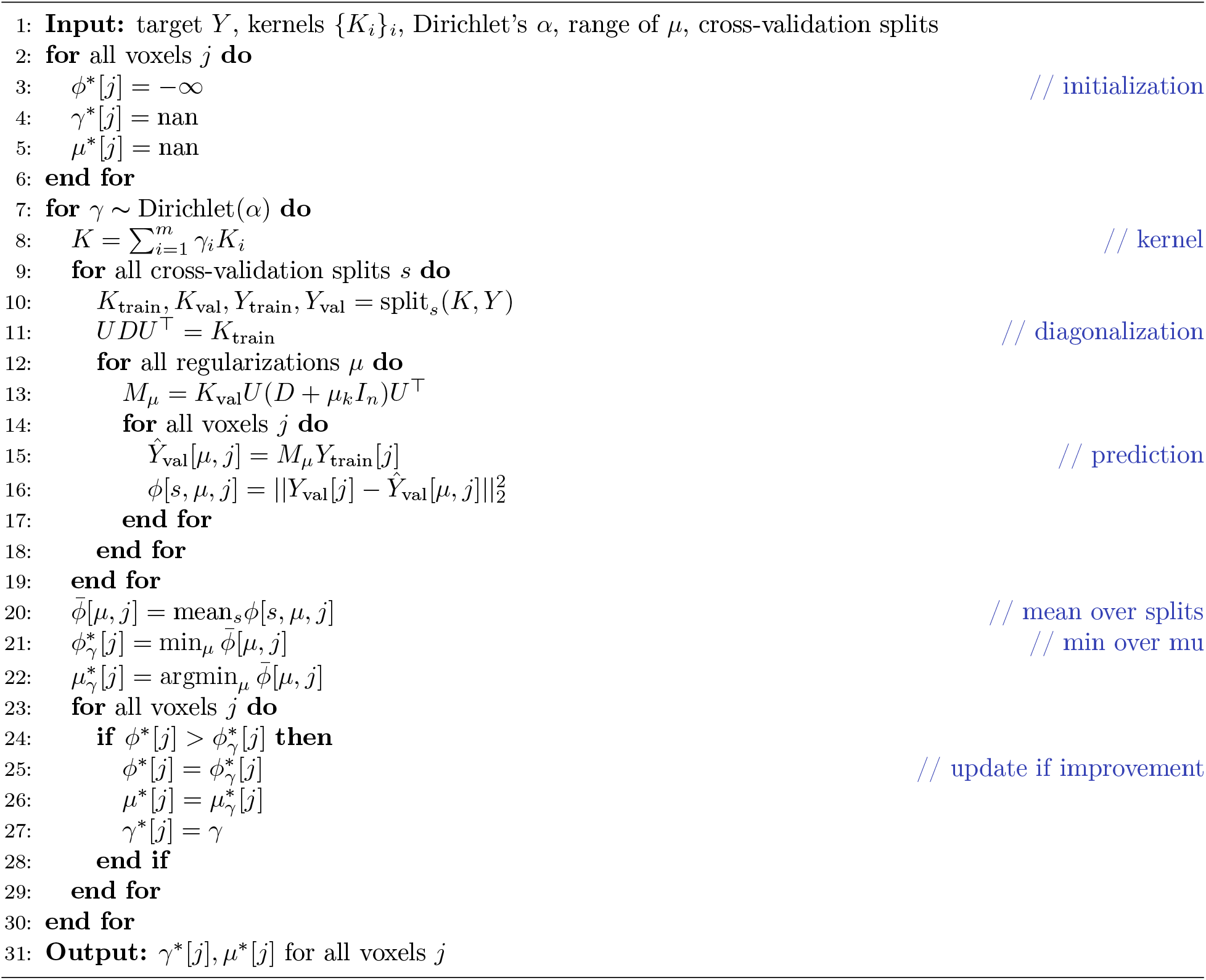

#### Algorithm 5

Multiple-kernel ridge regression solver - Hyperparameter gradient descent

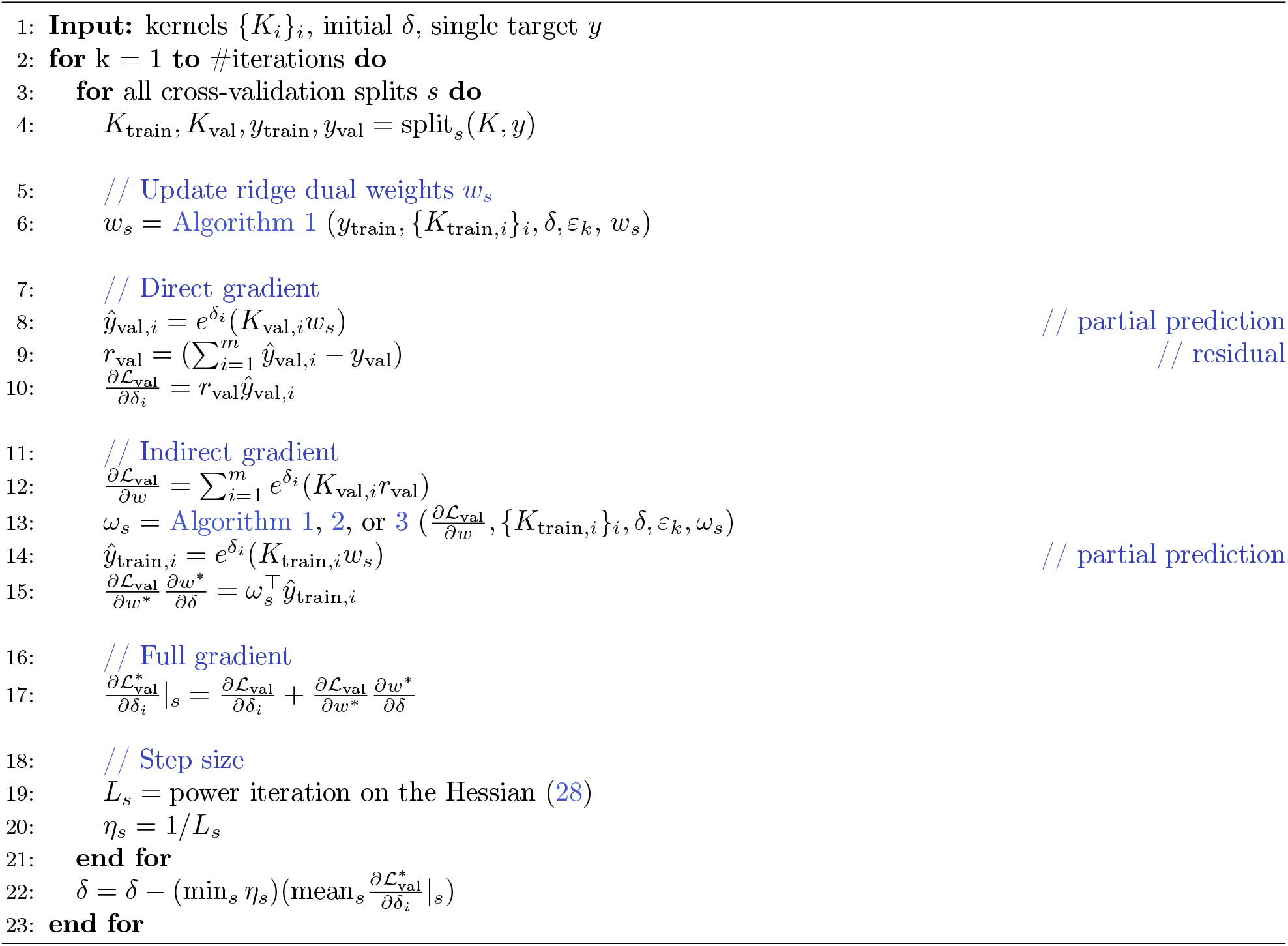

## Footnotes

1 https://github.com/gallantlab/himalaya

2 https://github.com/gallantlab/tikreg

3 https://github.com/gallantlab/himalaya

## Notes

https://github.com/gallantlab/himalaya

### Competing Interest Statement

The authors have declared no competing interest.

### Summary of Updates

Added missing author

